# Super-silencer perturbation by EZH2 and REST inhibition leads to large loss of chromatin interactions and reduction in cancer growth

**DOI:** 10.1101/2023.08.29.555291

**Authors:** Ying Zhang, Kaijing Chen, Seng Chuan Tang, Yichao Cai, Akiko Nambu, Yi Xiang See, Chaoyu Fu, Anandhkumar Raju, Benjamin Lebeau, Zixun Ling, Marek Mutwil, Manikandan Lakshmanan, Motomi Osato, Vinay Tergaonkar, Melissa Jane Fullwood

**Author notes:** Correspondence should be sent to: Melissa J. Fullwood, School of Biological Sciences, Nanyang Technological University. These authors contributed equally (Ying Zhang, Kaijing Chen, Seng Chuan Tang, Yichao Cai).

## Abstract

Human silencers have been shown to exist and regulate developmental gene expression. However, the functional importance of human silencers needs to be elucidated, such as whether they can form “super-silencers” and whether they are linked to cancer progression. Here, through interrogating two putative silencer components of *FGF18* gene, we found that two nearby silencers can cooperate via compensatory chromatin interactions to form a “super-silencer”. Furthermore, double knockout of two silencers exhibited synergistic upregulation of *FGF18* expression and changes of cell identity. To perturb the “super-silencers”, we applied combinational treatment of an EZH2 inhibitor GSK343, and a REST inhibitor, X5050 (“GR”). We found that GR led to severe loss of TADs and loops, while the use of one inhibitor by itself only showed mild changes. Such changes in TADs and loops were associated with reduced CTCF and TOP2A mRNA levels. Moreover, GSK343 and X5050 synergistically upregulated super-silencer-controlled genes related to cell cycle, apoptosis and DNA damage, leading to anticancer effects both *in vitro* and *in vivo*. Overall, our data demonstrated the first example of a “super-silencer” and showed that combinational usage of GSK343 and X5050 to disrupt “super-silencers” could potentially lead to cancer ablation.

## Introduction

Cis-regulatory elements are important for controlling gene expression including active elements such as enhancers and super-enhancers (SEs)^1,2^, and repressive elements such as silencers^3-6^. SEs are stitched together by multiple enhancers and bound by master transcription factors such as OCT4 and SOX2 to drive expression of genes that define cell identity^2^. Different components of SE can work in different modes to activate target gene expression: hierarchical^7,8^, additive^9,10^, and redundant^11,12^.

SEs can be acquired by key oncogenes (such as the *MYC* oncogene in cancer) to drive the process of tumorigenesis^13^. SEs are highly associated with chromatin interactions^14^, and can connect with the target oncogene via long-range chromatin interactions^15,16^. Altered chromatin interactions have been observed to drive the expression of oncogenes such as *TERT*^17^. Therefore, inhibition of cancer-specific SEs became a new direction to investigate for cancer therapies^16^. Pharmacological inhibition of transcriptional activators that are involved in SE function can be one approach to treatment of cancer^18^. For example, inhibitors of BET bromodomain proteins, which are enriched in SEs, are now being used in clinical trials and show therapeutic potential across multiple cancer types^19^. The disruption of long-range chromatin interactions between SEs and target oncogenes could be another strategy used to disrupt the SEs. Recently, one class of anti-cancer drugs, curaxins^20^, has been found to disrupt long-range chromatin interactions both *in vitro* and *in vivo*, thus suppressing the expression of key oncogenes, especially *MYC* family genes^21^.

Silencers, which are regions of the genome that are capable of repressing gene expression, can be predicted by various methods based on histone marks and chromatin interactions^3-6^. Different methods have been proposed to identify silencers in human and mouse genomes, including correlation between H3K27me3-DNaseI hypersensitive site and gene expression^6^, subtractive approach^3^, and PRC2 Chromatin Interaction Analysis with Paired-End Tag sequencing (ChIA-PET)^4^. Our previous work demonstrated a method of identifying silencers through ranking and stitching the histone H3K27me3 peaks, through which to identify H3K27me3-rich regions (MRRs) as human silencers^5^. We validated the gene silencing function of selected MRRs and showed that they can control genes related to cell identity^5^, which is in line with several other studies^3-6^. We also found that MRRs were highly associated with chromatin interactions and validated two looping silencers connecting to silenced genes. These results showed that MRRs exhibit cell-type specificity and enrichment for chromatin interactions, which is consistent with the characteristics of SEs. Therefore, we speculated that at least some MRRs could function as “super-silencers”. Here a “super-silencer” (SS) is defined as a genomic region of high H3K27me3 signal comprised of multiple silencers that can work as a single entity to repress transcription of genes that are important to the regulation of cell identity.

Studies of enhancers have revealed their impact on specificity and robustness of transcriptional regulation and their importance during development and evolution^2,13,22,23^. Moreover, epigenetic drugs that target SEs have shown good efficiency in terms of killing cancer cells^18^. Similarly, it is crucial to address the mechanism of action of silencers, especially how they may work in tandem as a SS to silence gene expression. Finding drugs that can target silencers or SS may also reveal another dimension for cancer therapies, similar to drugs that target SEs.

Here, we demonstrated that two silencer components inside one MRR can cooperate as a SS through compensated chromatin interactions to synergistically repress the *FGF18* gene. The 3D genome organization of silencer components underlies such synergism. We disrupted the SS and their associated chromatin interactions with the epigenetic drugs GSK343 and X5050 targeting EZH2 and REST, respectively. We were surprised to find that a single treatment of either GSK343 or X5050 mildly changed the genome organization and cancer cell viability, while combined treatment with GSK343 and X5050 (“GR”) exerted synergistic effects. GR also showed antitumor effects, suggesting that the usage of combined treatment to target SS and 3D genome organization should be further explored as a potential future cancer therapy.

## Results

### Removal of two silencers in the same MRR leads to synergistic upregulation of *FGF18* expression and growth inhibition

In our previous work, we experimentally validated two powerful distal silencers in the human genome: the silencer loops to *IGF2* gene and *FGF18* gene, respectively^5^. To investigate the working mode of silencer components of a SS, we further dissected the two component silencers (silencer1 and silencer2, termed “S1” and “S2”) within the same MRR, which are distal to the *FGF18* gene. We generated individual CRISPR knockouts (KO) for each component silencer (S1KO and S2KO) and a combinatorial double KO (DKO) (Figure 1A, Figure S1A-B).

**Figure 1.**
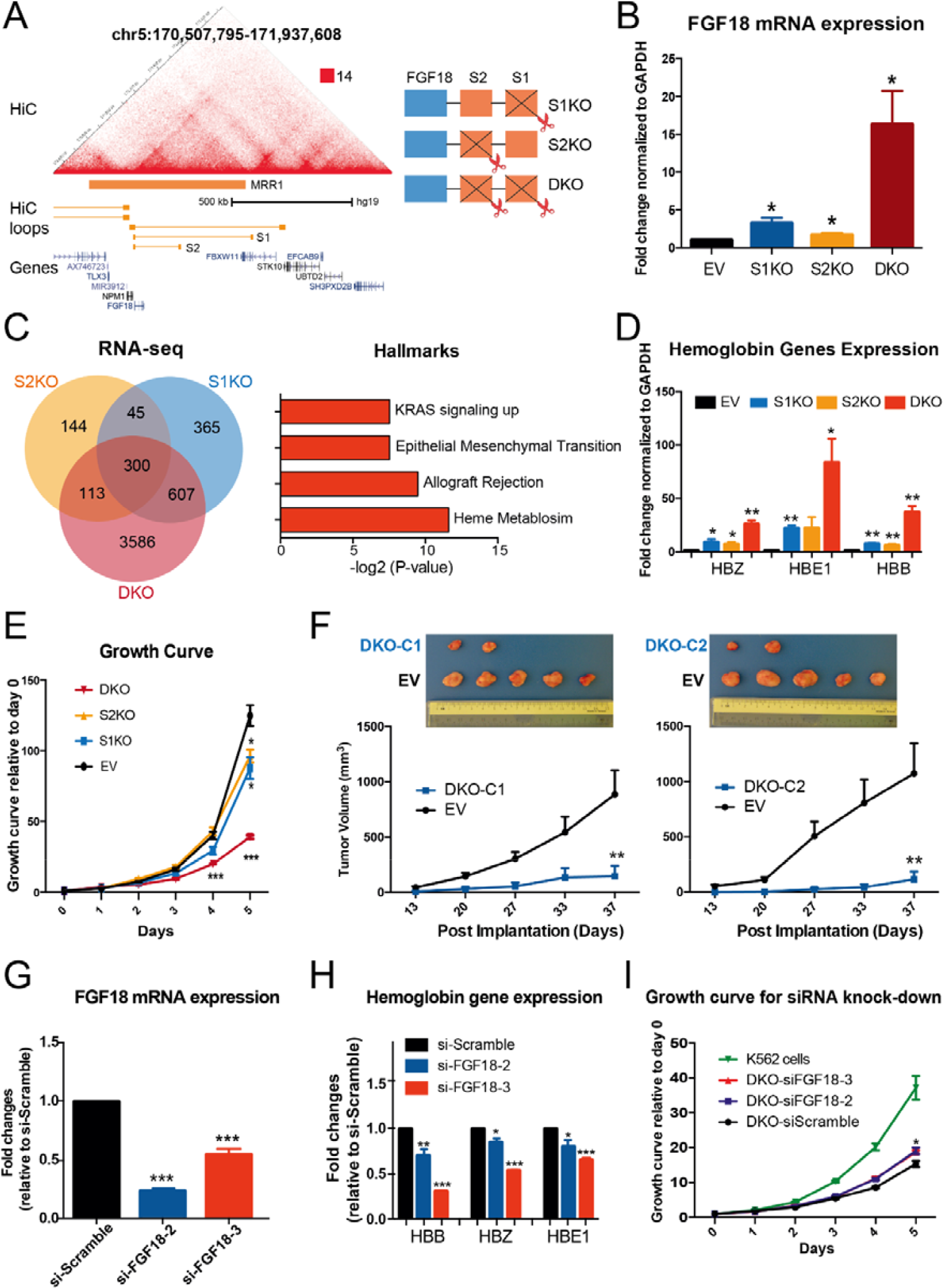
Two silencers removal leads to synergistic upregulation of *FGF18* expression and tumor inhibition. **A**. Hi-C matrix and loops at MRR1 region showing two looping silencers (S1 and S2) that exhibit chromatin interactions with *FGF18* gene in K562 cells^57^. Schematic depicting strategy to delete different silencer components to generate S1 knockout cells (S1KO), S2 knockout cells (S2KO) and double knockouts of S1 and S2 (DKO) in K562 cells using CRISPR/Cas9 gene-editing tool. **B**. RT-qPCR analysis of expression of *FGF18* in vector control clone (“Empty Vector”; “EV”), S1KO cells, S2KO cells, and DKO cells. Relative fold change normalized to GAPDH plotted. **C**. Venn diagram depicting number of significant differentially expressed genes in RNA-seq data of S1KO, S2KO and DKO cells as compared to EV cells. Gene Set Enrichment Analysis (GSEA) performed using intersection of significant differentially expressed genes in different RNA-seq (300 genes) against cancer hallmark database^58^. Data shown as –log_2_(p value). **D**. RT-qPCR analysis of expression of haemoglobin genes (*HBZ, HBE1* and *HBB*) in EV, S1KO, S2KO and DKO cells. Fold change normalized to GAPDH plotted. **E**. Growth curves in EV cells and different knockout cells. Data calculated as fold change against day 0. **F**. Representative tumors pictured at final day (top) and tumor volume (mm^3^) with different post implantation days (bottom) shown. Tumor growth in SCID (Severe Combined Immunodeficiency) mice injected with two different DKO clones (DKO-C1 and D1KO-C2) and EV cells. N=5 for each group. **G and H**. RT-qPCR analysis of expression of *FGF18* (G) and hemoglobin genes (*HBZ, HBE1* and *HBB*) (H) upon small interfering RNA (siRNA) knockdown in DKO clone. Knockdown experiment is performed using scrambled oligonucleotides (siScramble) and two different siRNAs targeting human *FGF18* (siFGF18). Data shown as fold change relative to siScramble. **I**. Growth curves of wild type K562 cells and DKO clones transfected either with siScramble or siFGF18. Data calculated as fold change against day 0. Data shown as average + standard error of mean (SEM). P value calculated using two-tailed student’s t-test. P value less than 0.05, 0.01 or 0.001 shown as *, ** or ***, respectively.

First, we checked the influence of our different KO clones on *FGF18* expression. As expected, S1KO and S2KO both showed *FGF18* expression upregulation compared to the empty vector (EV) (Figure 1B, Figure S1C), suggesting that both S1 and S2 can function as silencers. However, we observed that each silencer exerted a different extent of gene upregulation upon removal, hinting that they may not have equivalent silencing capabilities. Remarkably, DKO showed an upregulation of FGF18 greater than the sum of S1KO and S2KO (Figure 1B, Figure S1C). This observed synergism between the two silencer components indicates that the MRR may behave as a SS.

Next, we wanted to know whether DKO confers synergistic effects in terms of phenotype. We previously observed an increase in cell adhesion ability in S1KO^5^, but this phenotype did not translate to the DKO cells (Figure S1D). To explore further phenotypes, we first performed RNA-seq in the different KO cell lines. Consistent with the effect on FGF18, DKO led to the most differentially expressed genes (DEGs) (Figure 1C, Figure S1E). Next, we overlapped the RNA-seq data to identify DEGs common to all three KO clones. Interestingly, genes commonly altered were significantly enriched for several cancer hallmarks, including pathways associated with the erythroid differentiation phenotype, such as heme metabolism (Figure 1C). As we previously elucidated this phenotype in S1KO cells^5,24^, we investigated if the DKO would potentiate it. To do so, we measured the expression of erythroid differentiation indicators (haemoglobin genes: *HBZ, HBE1* and *HBB*)^25^, as we did previously^5^. We found that DKO dramatically increased these erythroid differentiation markers compared to single KO clones (Figure 1D, Figure S1F), consistent with the synergistic upregulation of *FGF18* expression (Figure 1B, Figure S1C).

Leukemic erythroid differentiation can cause cell growth inhibition, which is one of the existing therapies for leukemia^26,27^. Therefore, we asked if DKO also shows synergistic effects in terms of growth inhibition. We performed cell growth assay *in vitro* for different KO clones (Figure 1E, Figure S1G) and xenograft experiments *in vivo* by injecting two different DKO clones into mice (Figure 1F). We found that DKO had synergistic growth inhibition *in vitro* and *in vivo*, which was in line with the synergistic erythroid differentiation phenotype of DKO.

To confirm that the phenotypes we observed were associated with the upregulation of *FGF18* gene expression, we performed siRNA knockdown experiments against the *FGF18* gene in DKO cells. Downregulation of FGF18 in DKO cells (Figure 1G) reduced the expression of erythroid differentiation markers (Figure 1H) and partially restored cell growth capabilities (Figure 1I). Similarly, we also overexpressed FGF18 in wild type K562 cells by adding recombinant human protein FGF18 (rhFGF18). The overexpression of FGF18 significantly inhibited cell growth (Figure S1H), although to a lesser extent than the DKO cells. Then, to determine if other genes near the FGF18 locus were affected by the deletion of S1 and/or S2, we performed qRT-PCR on RNA isolated from EV and our knockout clones for five genes located within 1Mb of the FGF18 locus. The expression of most genes was either unaffected or minimally affected by the single or double removal of S1 and/or S2 (Figure S1I). Therefore, these experiments support a direct and specific relationship between *FGF18* expression and the observed phenotypes.

In sum, the knockout of both silencers resulted in the upregulation of FGF18 which partially drove changes in gene expression characteristic of erythroid differentiation and inhibition of cell growth. Taken together, our results demonstrate that two silencer components can act synergistically, as a super-silencer, to repress the expression of genes crucial to maintenance of cancer cell identity, such as FGF18, and influence biological phenotypes. This highlights the potential of super-silencers in oncological studies.

### 3D genome organization underlies the synergism of silencers

Next, we aimed to detect if the synergy observed between S1 and S2 in regulating gene expression would translate to the physical interactions between S1 and S2 and chromatin organization changes in different KO cells. Therefore, we first performed 4C-seq using *FGF18* promoter as the viewpoint in EV, S1KO and DKO clones to investigate whether chromatin interactions may underlie gene expression modulation by distant silencers^15,28-30^.

Interestingly, we found that silencers do participate in the maintenance of the chromatin interaction landscape. Indeed, S1KO led to 24 gained chromatin loops and 3 lost chromatin loops, connecting *FGF18* to more distant loci, compared to control cells (Figure 2A). Conversely, DKO cells lost 10 chromatin loops compared to control cells and gained 17 chromatin loops. In contrast with the loops gained by S1KO cells, DKO gained loops were predominantly within the region between *FGF18* and S2. The chromatin loops gained by DKO cells also tended to be shorter, further constraining all *FGF18* chromatin loops to a narrow region between FGF18 gene and S2, which noticeably changed the local chromatin interaction landscape (Figure 2B). Consistent with our previous observations^5^, S1KO and DKO both showed that long distance loops tended to be gained and lost more easily than nearby loops (Figure S2A-B).

**Figure 2.**
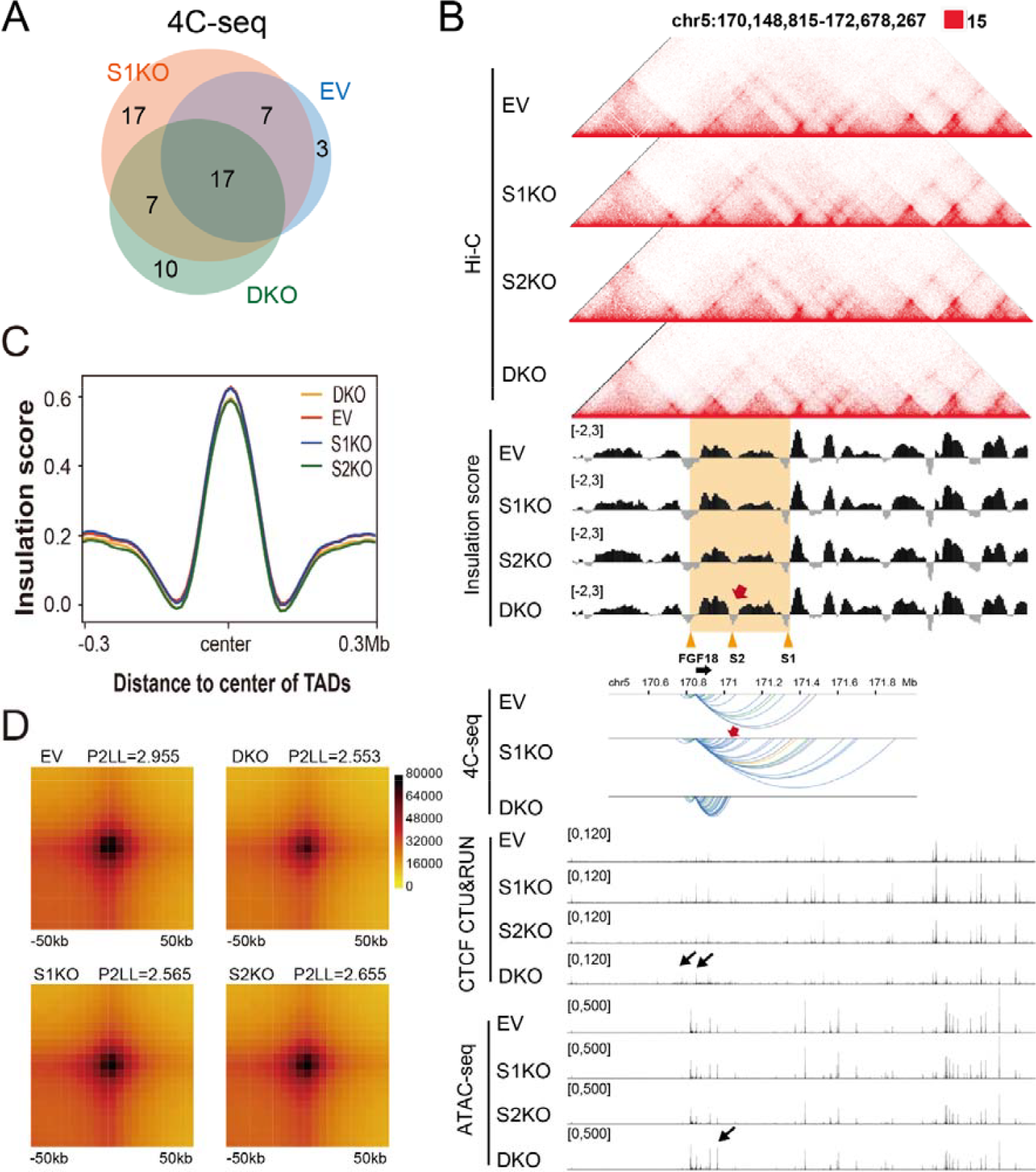
3D genome organization underlies synergism of super-silencers. **A**. Venn diagram depicting significant chromatin interactions identified by 4C-seq using *FGF18* promoter as viewpoint in EV, S1KO and DKO cells. Two replicates of 4C-seq were performed and significant chromatin interactions (p<0.05) were analyzed by R3Cseq package^59^. **B**. Screenshot depicting aligned Hi-C matrix, insulation score, CTCF CUT & RUN and ATAC-seq signal tracks at *FGF18* SS region in EV, S1KO, S2KO and DKO cells, as well as 4C-seq of EV, S1KO and DKO at *FGF18* SS region. *FGF18* SS region is highlighted by orange box and some specific sites (*FGF18* gene region, S2 and S1) are indicated. Increased ATAC-seq peaks and Cut & Run peaks are indicated by black arrows. Colors of arcs represent histone states of chromatin interactions (yellow: H3K27ac-associated loops; blue: H3K27me3-associated loops; green: both H3K27ac- and H3K27me3-associated loops; grey: no H3K27ac- and H3K27me3-associated loops)^5^. Red arrows indicate new insulation peak at S2 site in DKO cells coupled with compensated chromatin interactions to S2 site in S1KO cells. **C**. Mean plot describing genome-wide insulation score around TADs (use TADs in EV as reference TADs) in EV and different KO cells (S1KO, S2KO and DKO). **D**. Aggregate peak analysis (APA) for all loops in EV, S1KO, S2KO and DKO cells (use EV loops as reference). Loops are aggregated at the center of a 50kb window at 5kb resolution. The ratios of signal at the peak signal enrichment (P) to the average signal at the lower left corner of the plot (LL) (P2LL) are indicated to show the normalized intensity of all loops.

Surprisingly, when we carefully dissected the loops in S1KO, we found multiple new loops connecting *FGF18* to the S2 site in S1KO cells (Figure 2B), suggesting that S2 can partially compensate for the role of S1 through increased contact frequency with *FGF18* in S1KO. This partial compensation may explain why knockout of the two silencers leads to synergistic effects.

To further characterize whether the TAD and loop changes occurred locally at the *FGF18* silencer region or genome-wide, we performed Hi-C in EV, S1KO, S2KO and DKO cells. We found that the number of TADs and loops were similar across different KO cells (Figure S2C). When inspecting the *FGF18* silencer region, we observed changes in the Hi-C matrices where the loops between *FGF18* gene promoter and S1 were lost and the sub-TAD between the *FGF18* gene promoter and S2 was stronger in DKO (Figure 2B). By calculating the insulation score, a new boundary was identified at the S2 site in the DKO cells, which again indicated the formation of a stronger sub-TAD formation between the *FGF18* gene promoter and the S2 site (Figure 2B, Figure S2D). This was also concordant with the 4C-seq data which showed that all the chromatin interactions in the DKO cells were constrained inside of this strong sub-TAD region (Figure 2B).

Apart from the *FGF18* silencer region, other genomic regions did not show any obvious changes. We explored genome-wide changes in TADs and loops by calculating the insulation score for all the TADs in different KO cells (Figure 2C). Data showed that the average insulation scores for all the TADs in different samples were highly similar (Figure 2C). We also analysed the loop strength by aggregate peak analysis (APA) plot for different KO cells (Figure 2D), which showed that the loop strength was also highly similar across all four samples (Figure 2D). The Hi-C matrix of another MRR region at the *IGF2* gene, which was previously confirmed to be a silencer region^5^, did not show any obvious changes (Figure S2E), suggesting that our KO of the silencers of *FGF18* did not affect the 3D genome organization of other MRR regions. Together with the insulation score analysis for the TADs and the APA plot, our results suggested that removing component silencers resulted in local changes specific to the *FGF18* region and did not lead to genome-wide changes. This is consistent with the notion of a SS that mainly regulates genes within its vicinity.

As CTCF binding is an important factor in the definition and insulation of TAD and sub-TADs boundaries^31^, we performed CTCF Cleavage Under Targets and Release Using Nuclease (Cut & Run) assay in EV and different KO cells^32^. Overall, we observed a modest genome-wide increase in CTCF binding sites in all KO cells as compared to EV (Figure S2F). More specifically, we also observed an increase in CTCF binding sites within the FGF18 SS in S1KO, S2KO and DKO cells, compared with EV cells (Figure S2F). Therefore, the reorganization of chromatin conformation driven by the DKO could be potentiated by the recruitment of additional CTCF binding sites.

Changes in 3D genome organization are often associated with altered chromatin accessibility^33^. Therefore, we investigated chromatin accessibility changes upon silencer loss using ATAC-seq in EV and different KO cells. After single and double KO of component silencers, we observed a modest increase in ATAC-seq open chromatin regions genome-wide and within the *FGF18* SS (Figure 2B, Figure S2G). However, chromatin accessibility around *FGF18* gene promoter and within the newly strengthened sub-TAD noticeably and specifically increased upon the combined loss of S1 and S2 (Figure 2B). These findings correlate with the synergistic upregulation of *FGF18* expression in DKO cells (Figure 1B).

Overall, we showed that the component silencers S1 and S2 work together as a SS, and that knocking out these component silencers destabilizes the local 3D genome organization. Furthermore, these organization changes are linked to recruitment of epigenetic regulator CTCF and altered chromatin accessibility associated with the transcriptional upregulation of the SS target genes.

### Epigenomic differences together with chromatin interactions underlie the action of the “super-silencer”

In addition to investigating 3D genome organization, we further explored the association between silencer synergism and changes in histone modifications, H3K27ac and H3K27me3, associated with increased and decreased of transcriptional activities, respectively^34,35^. To do so, we performed H3K27ac and H3K27me3 ChIP-seq in EV, S1KO and DKO cells. At transcription start sites (TSS) genome-wide, H3K27ac levels were not consistently altered by S1KO or DKO compared to EV (Figure S3A). However, when looking specifically at the FGF18 locus in DKO cells, H3K27ac displayed a marked increase in signals at multiple peaks, compared to S1KO and EV, which we validated by ChIP-qPCR (Figure S3B). On the other hand, S1KO and DKO resulted in a noticeable decrease in the H3K27me3 signal at TSS genome-wide (Figure S3A), while the changes at the FGF18 locus between DKO and EV were moderate compared to H3K27ac (Figure S3B). Overall, this reprogramming of histone marks is consistent with the pronounced upregulation of FGF18 in DKO compared to S1KO and EV. Furthermore, we aimed to associate the changes in H3K27ac and 3D genome organization at this locus. Therefore, we performed an integrative analysis of 4C-seq and ChIP-seq comparing EV vs DKO, EV vs S1KO and S1KO vs DKO (Supplementary Text). We found that DKO chromatin loops were associated with increased H3K27ac ChIP-seq signals compared to both EV and S1KO chromatin loops, but no similar increase in H3K27ac ChIP-seq signal was observed between S1KO and EV chromatin loops, possibly due to the compensation by S2 (Figure S3C-H).

The increased H3K27ac signals in DKO gained loops suggested that there might be new enhancers that looped to the *FGF18* gene promoter in DKO. Indeed, we identified two putative enhancers gained in DKO cells (Figure S3I-J). Using 3C-PCR and Sanger Sequencing, we validated the presence of a chromatin loop between the gained enhancer proximal to FGF18 (Figure S3I). Taken together, our results showed that silencer knockout leads to significant changes in both chromatin interactions and histone modifications in DKO cells, explaining the synergistic upregulation of *FGF18*.

This work is, to the best of our knowledge, the first indication that two distal silencers can cooperate to function as a SS. Also, disease-associated DNA sequence variation has been shown to be enriched in regulatory regions, such as SEs^13^. Therefore, we reasoned that SS could also be enriched and help to identify key genes that need to be kept silenced for maintenance of cancer phenotypes. We performed a similar analysis of disease-associated DNA sequences and showed that compared to the random regions, SS were enriched for SNPs associated with the groups: all diseases, all cancers, Acute Myeloid Leukemia (AML) and Chronic Myeloid Leukemia (CML) (Figure S3K-L). This analysis highlights the potential involvement of SS in diverse pathologies, including cancer. Therefore, SS could be therapeutic targets for treating cancer.

### Combinational treatment of GSK343 and X5050 leads to synergistic loss of TAD and loops

Since our putative SS are enriched for H3K27me3 histone modifications, we asked whether depletion of H3K27me3 affects the 3D genome organization of SS. We treated wild type K562 cells with a EZH2 histone methyltransferase inhibitor, GSK343 (5 µM), to deplete H3K27me3 histone modifications. We performed a genome-wide Hi-C analysis and observed modest changes in and around 10% of the genome-wide TADs and loops upon GSK343 treatment (Figure 3A-B, Figure S4A-B).

**Figure 3.**
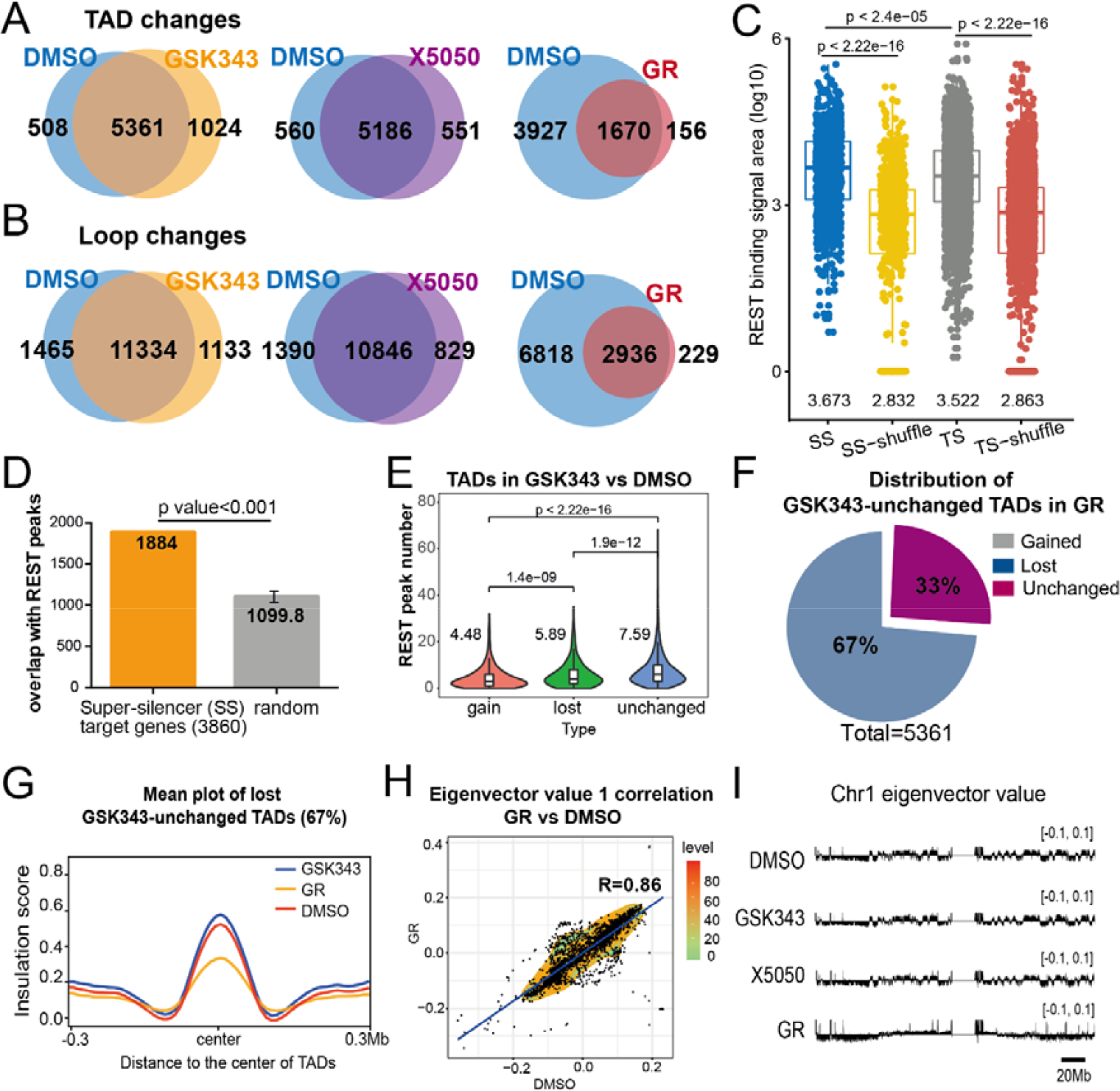
Combinational treatment of GSK343 and X5050 leads to synergistic losses of TADs and loops. **A and B**. Venn diagrams of TAD changes (**A**) and loop changes (**B**) in DMSO, GSK343, X5050 and combinational treatment of GSK343 and X5050 (“GR”) in K562 cells. **C**. Boxplot depicting REST enrichment at SS and typical silencers (TS) in K562 cells. SS-shuffle and TS-shuffle serve as random control. REST enrichment shown as REST binding signal area (log10). **D**. Bar chart showing REST enrichment at SS target genes and random genomic regions in K562 cells. Y-axis shows number of overlaps between SS target genes/random genomic regions and REST ChIP-seq peaks. **E**. Violin plot depicting number of REST peaks at different TAD categories (gained, lost and unchanged) in GSK343 vs DMSO condition in K562 cells. **F**. Pie chart showing distribution of GSK343-unchanged TADs compared to TAD changes in GR vs DMSO condition. **G**. Mean plot depicting genome-wide insulation score around TADs of GSK343-unchanged TADs (67%) lost in GR vs DMSO condition. **H**. Density plot describing global correlation between eigenvector value from DMSO condition and GR condition at 1Mb resolution. X-axis represents eigenvector value in the DMSO condition, while Y-axis represents eigenvector value in GR condition at the same locus. **I**. Representative eigenvector value for 50kb resolution at chromosome 1 in DMSO, GSK343, X5050 and GR conditions.

We reasoned that just as SEs are associated with multiple transcription factors^13^, SSs might also be associated with multiple transcription factors. Therefore, a combination of epigenetic drugs targeting different repressive transcription factors would be needed to affect chromatin interactions at SS. As a first step in identifying additional targets, we aimed to identify potential SS regulated genes using H3K27me3 HiChIP. We identified 3860 genes associated with two or more H3K27me3 HiChIP loops, similar to *FGF18* gene, and categorized them as potential SS target genes. Next, we examined these SS and SS target genes for associated transcription factors. We found that REST, which is known to be a repressor of transcription^36^, was enriched at SS compared to random regions and typical silencers (Figure 3C). REST was enriched at the identified potential SS target genes as well (Figure 3D). Interestingly, REST was also enriched at TADs that remained unchanged after GSK343 treatment, compared to gained and lost TADs (Figure 3E), suggesting that REST may help to retain the structure of TADs upon GSK343 treatment. As such, we reasoned that combining a REST inhibitor with GSK343 may disrupt the structures of TADs and loops unaffected by single inhibition of H3K27me3.

We tested whether REST is directly involved in TAD formation and SS functioning. To do so, we applied a REST inhibitor, X5050, to K562 cells and performed Hi-C. Similar to the GSK343 treatment, the overall TAD and loop changes following X5050 treatment alone were modest (Figure 3A-B, Figure S4C-D). As for H3K27me3 inhibition, targeting REST alone cannot, consequently, ablate TAD structures. We first demonstrated a reduction in H3K27me3 signals at TSS genome-wide and REST protein levels in cells treated with both GSK343 and X5050 (referred to as “GR”), in comparison to the control cells (Figure S4E-F). Next, we tested if combined H3K27me3 and REST inhibition had an additive or synergistic effect on chromatin organization and SS. In contrast to single treatments, GR treatment caused severe loss of TADs and loops (Figure 3A-B, Figure S4G-H). Compared to DMSO, GR lost 3927 TADs and 6818 loops (Figure 3A-B). Among TADs that were unaffected by a single GSK343 treatment, 67% of them were lost following GR treatment (Figure 3F), together with decreased TAD insulation score (Figure 3G). These results denote the additive, potentially synergistic and combinatorial effect of H3K27me3 and REST inhibition, hinted at by the enrichment of REST binding at unchanged TADs following GSK343 treatment.

Additionally, besides the loss of TADs and loops, GR also showed altered A/B compartments (Figure 3H-I). On the other hand, GSK343 and X5050 treatment alone did not show obvious changes in A/B compartments (Figure S4I-J). We also compared the genome-wide changes in chromatin accessibility caused by GR with those detected following the double KO of silencers surrounding FGF18 from our previous model. As expected, the majority of gained and lost ATAC-Seq peaks following GR treatment were not similarly altered by the DKO. By contrast, the near totality of peaks unaltered by GR were also unchanged by DKO (Figure S4K). This analysis supports a genome-wide effect of GR, compared to the localized effect around FGF18 of DKO. These results further suggest that there are potential transcriptional alterations in the GR condition.

### GR treatment leads to apoptosis and cell cycle arrest through the upregulation of super-silencer controlled genes

To survey how GR treatment might affect gene expression, we first focused on its potential effects on SSs and SS-regulated genes. Since H3K27me3 and REST are both associated with SSs, we suspected that their inhibition would affect the chromatin organization at these sites disproportionately. Indeed, we observed that MRRs (putative SS sites) were enriched at lost TADs compared to a set of randomized equivalent regions (Figure 4A).

**Figure 4.**
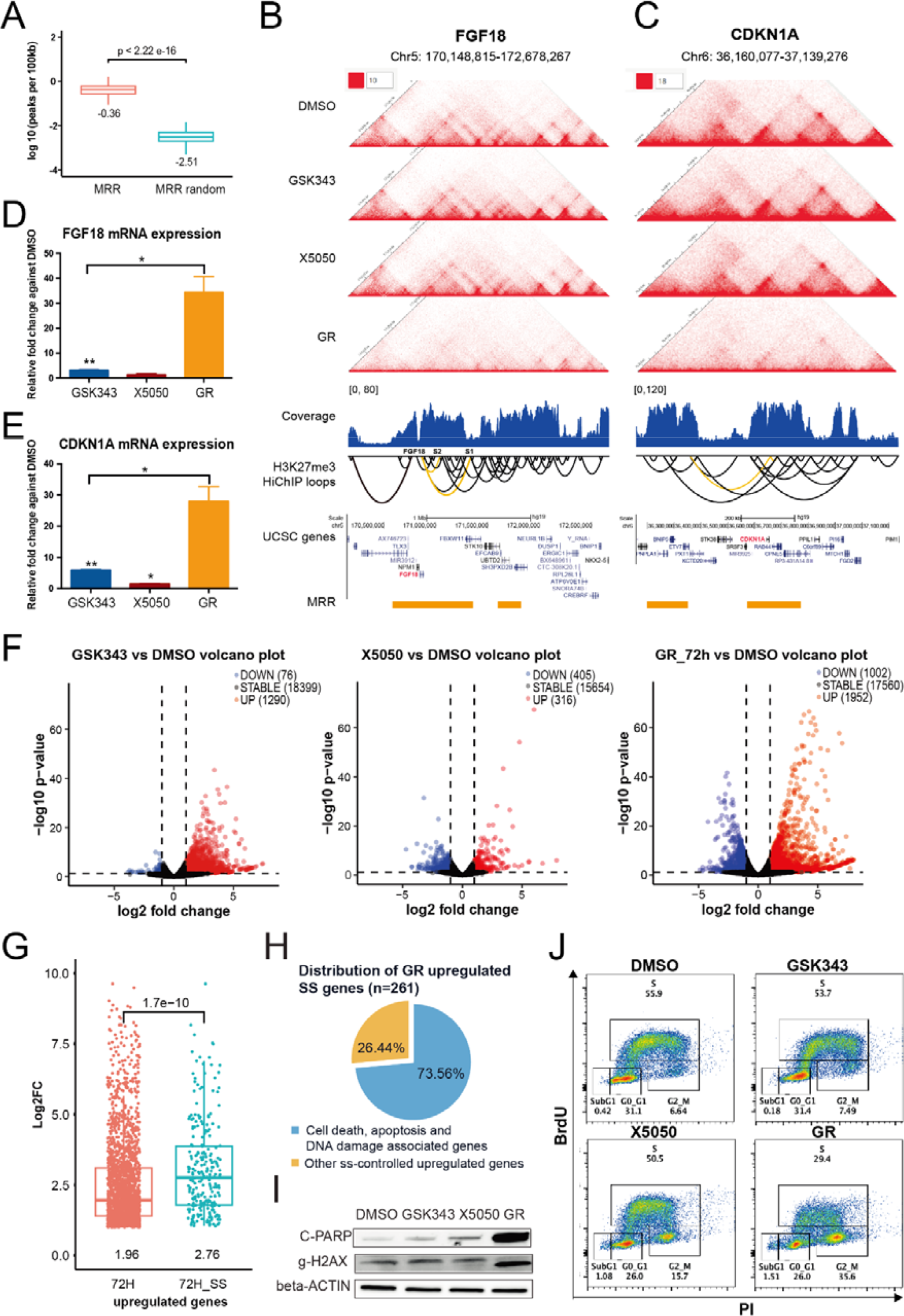
GR treatment leads to apoptosis and cell cycle arrest through upregulation of super-silencer controlled genes. **A**. Boxplot describing number of MRR peaks and randomly generated artificial MRR peaks per 100kb (log10 scale) on lost TADs (GR vs DMSO), along with corresponding p-value. **B**. Screenshot showing Hi-C contact matrix of *FGF18* region in DMSO, GSK343, X5050, and GR conditions. H3K27me3 HiChIP data including coverage and loops in normal K562 cells, UCSC gene tracks and MRR annotations are shown. H3K27me3 HiChIP loops associated with *FGF18* gene, S1 and S2 are indicated by yellow color. **C**. Screenshot showing Hi-C contact matrix of *CDKN1A* region in DMSO, GSK343, X5050, and GR conditions. H3K27me3 HiChIP data including coverage and loops in normal K562 cells, UCSC gene tracks and MRR annotations. H3K27me3 HiChIP loops associated with *CDKN1A* gene are indicated by yellow color. **D**. RT-qPCR analysis of expression of *FGF18* in GSK343, X5050 and GR conditions. Results shown as relative fold change against DMSO control. **E**. RT-qPCR analysis of expression of *CDKN1A* in GSK343, X5050 and GR conditions. Results shown as relative fold change against DMSO control. **F**. Volcano plots of RNA-seq comparing DMSO to GSK343 72 h, X5050 72 h, and GR 72 h conditions. The number of downregulated, stable and upregulated genes are indicated. **G**. Boxplot showing mRNA levels of upregulated genes and SS-controlled upregulated genes at 72 h following GR treatment. **H**. Pie chart showing distribution of GR upregulated SS-controlled genes according to their associations with apoptosis, DNA damage and cell cycle arrest pathways. **I**. Western blot showing protein levels of cleaved-PARP (C-PARP), gamma-H2AX (g-H2AX) and beta-actin in K562 cells treated with DMSO, GSK343, X5050 and GR for 72 h. **J**. Cell cycle analysis of K562 cells after exposure to DMSO, GSK343, X5050 and GR for 72 h. Cell cycle analysis is performed on flow cytometer. Percentage of subG1, G0/G1, S and G2/M phases indicated for different drug-treated conditions. Data shown as average + SEM. P value calculated using two-tailed student’s t-test. P value less than 0.05 or 0.01 shown as * or **, respectively.

As SSs are associated with gene-silencing interactions, their loss should result in the upregulation of their target genes. Therefore, we initially investigated gene expression changes for *FGF18*, previously established as a SS-controlled gene, and *CDKN1A*, another potential SS-regulated gene according to our H3K27me3 HiChIP and MRR mapping. Additionally, our Hi-C investigations revealed that both *FGF18* and *CDKN1A* displayed a clear loss of local interactions, overlapping with H3K27me3 regions, following GR treatment (Figure 4B-C). Similarly to when the SSs surrounding *FGF18* were deleted, GR treatment resulted in the upregulation of *FGF18* mRNA expression, detected by RT-qPCR (Figure 4D). *CDKN1A* mRNA levels also displayed clear upregulation (Figure 4E), supporting the repressive influence of SS interactions on transcriptional regulation, which are inhibited by GR treatment.

To investigate whether these observations were applicable genome-wide, we performed RNA-seq following DMSO, GSK343, X5050, and GR combinational treatment. Akin to the changes in chromatin conformation, transcriptional changes versus DMSO control were distinctly amplified in GR compared to single treatments (Figure 4F). Further, we observed that the GR combination led to almost twice as many genes being upregulated (1952) compared to downregulated (1002). Additionally, the mRNA levels of SS related genes were significantly higher compared to all upregulated genes (Figure 4G). These results are consistent with a global loss of SS related TADs and loops in GR treatment, which repress gene expression in control condition and lead to a disproportionate transcriptional upregulation when inhibited.

Next, we aimed to determine if the upregulation of SS-related genes following the combined inhibition of H3K27me3 and REST were associated with specific functions or pathways. Using KEGG pathway analysis of upregulated genes following GR treatment, we observed that pathways related to apoptosis, cell cycle and DNA damage were significantly upregulated (Figure S5A). A similar trend was observed when looking specifically at SS upregulated genes, as 74% of them were part of either apoptosis, cell cycle, or DNA damage pathways (Figure 4H, Figure S5B). Therefore, we decided to validate whether these gene expression changes were driving biological phenotypes. We surveyed the protein expression of cleaved-PARP, a marker of apoptosis^37^, and γH2AX, a marker of DNA damage^38^, and found that they were uniquely increased by GR treatment (Figure 4I, Figure S5C). This was further supported by a disproportionate increase in total apoptotic cells at 72 h following GR treatment (Figure S5D). Similarly, flow cytometry analysis of the cell cycle showed a pronounced increase in the population of G2/M stage cells following GR treatment, compared to individual inhibition of H3K27me3 or REST (Figure 4J), denoting potential cell cycle arrest.

These results suggest that the global upregulation of SS-related genes associated with the loss of TADs and loops caused by GR treatment promotes apoptosis and cell cycle arrest in K562 leukaemia cells.

### Gradual depletion of CTCF and TOP2A explains the kinetics and amplitude of the loss of TADs and loops caused by GR treatment

Compounds targeting chromatin or epigenetic effectors can cause swift changes in 3D conformation, such as inhibitors targeting the Facilitates Chromatin Transcription (FACT) complex (curaxins)^20,21^, BET1 bromodomain inhibitors (JQ1)^39^, HDAC inhibitors (Trichostatin A and suberoylanilide hydroxamic acid)^40^, and RNA polymerase I and II-dependent transcription inhibitors (triptolide)^41^. To gain a better understanding of GR epigenetic effects driving the loss of loops and TADs we observed following GR treatment, we investigated the kinetics of the chromatin conformation changes.

To study the kinetics of 3D genome organization changes, we used Hi-C after 8 h and 24 h of GR treatment, in addition to the DMSO and 72 h previously described. Interestingly, the global losses of loops and TADs was observed only at 72 h, as denoted by a decrease in the number of TADs and loops called after 72 h compared with other time points (Figure S6A) and the marked reduction in TADs insulation score (Figure 5A) and aggregate peak intensity (Figure 5B). These observations suggest that GR treatment causes changes in 3D genome organization through a gradual or secondary mechanism, such as changes in gene expression.

**Figure 5.**
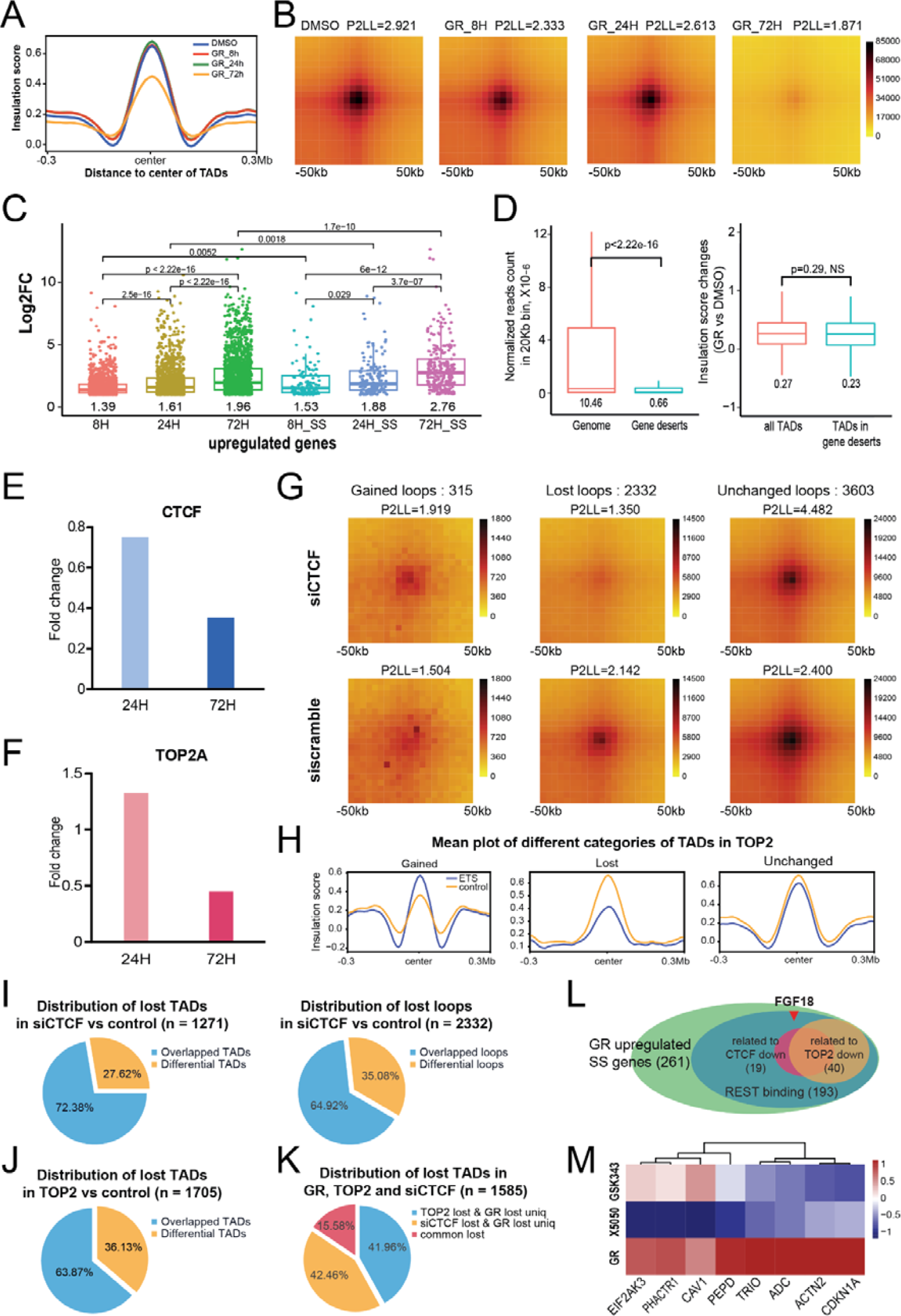
Gradual depletion of CTCF and TOP2A explains kinetics and amplitude of losses of TADs and loops caused by GR treatment. **A**. Mean plot describing genome-wide insulation score around TADs (use TADs in DMSO as reference TADs) in DMSO- and GR-treated cells at different time points (GR_8h, GR_24h and GR_72h). **B**. APA for all loops (use loops in DMSO as reference) in DMSO- and GR-treated cells at different time-points (GR_8h, GR_24h and GR_72h). Loops are aggregated at the center of a 50kb window at 5kb resolution. The ratios of signal at the peak signal enrichment (P) to the average signal at the lower left corner of the plot (LL) (P2LL) are indicated to show the normalized intensity of all loops. **C**. Boxplot showing genes and SS-controlled genes upregulated upon GR treatment at 8 h, 24 h and 72 h. **D**. Boxplot describing RNA normalized read count in 20 kb-long bin calculated from DMSO-treated K562 RNA-seq (left). P value indicated for these two categories. Boxplot describing insulation score changes (GR vs DMSO) for either all TADs or TADs located in gene deserts (right). P value indicated for comparison and NS stands for no significance. **E**. RNA expression of *CTCF* in GR-treated K562 cells at 24 h and 72 h as compared to GR at 8 h. **F**. RNA expression of *TOP2A* in GR-treated K562 cells at 24 h and 72 h as compared to GR at 8 h. **G**. APA for gained loops (left), lost loops (middle) and unchanged loops (right) in siCTCF (top) and siScramble (bottom). Loops are aggregated at the center of a 50 kb window at 5 kb resolution. The ratios of signal at the peak signal enrichment (P) to the average signal at the lower left corner of the plot (LL) (P2LL) are indicated to show the normalized intensity of all loops. **H**. Mean plots showing genome-wide insulation score around TADs of gained TADs, lost TADs, and unchanged TADs in DMSO control cells and etoposide-treated K562 cells. **I**. Pie charts showing distribution of overlapped and differential TADs (left) or overlapped and differential loops (right) lost upon siCTCF or GR treatments comparing to siScramble or DMSO control, respectively. **J**. Pie chart showing distribution of overlapped and differential TADs lost upon TOP2 inhibitor (etoposide) or GR treatments comparing to siScramble or DMSO control, respectively. **K**. Pie chart showing distribution of TADs that are commonly lost in siCTCF or etoposide compared to lost TADs in GR condition. **L**. Schematic describing large overlap of REST binding genes with GR upregulated SS-controlled genes, for which there is a large association with decreased mRNA expression of CTCF and TOP2. The number of genes is indicated in each category. *FGF18* is one of 193 genes identified in category with GR upregulated SS-controlled genes with REST binding. **M**. Heatmap showing gene expression of eight shortlisted REST binding-GR upregulated SS genes, in which these genes are also associated with lost TADs upon GR treatment and mRNA downregulation of CTCF and TOP2. Gene expression shown by log2 fold change in GSK343, X5050, and GR against DMSO condition.

To study the kinetic interplay of changes in gene expression and 3D genome organization, we also performed RNA-Seq after 8 h and 24 h of GR treatment, to the previously described DMSO and 72 h. Contrarily to the 72 h GR treatment previously shown, 8 h and 24 h treatments did not lead to a disproportionate upregulation of gene expression (Figure S6B). Interestingly, compared to the significant, but modest, gradual increase in gene upregulation from 8 h to 72 h, SS-related upregulated genes displayed a significantly more pronounced increase (Figure 5C). Therefore, although gene expression was altered early on following GR treatment, the upregulation of SS-related genes was mostly observed at 72 h, when loops and TADs loss became prominent.

Since active transcriptional activity can influence nearby 3D genome organization^41^, we aimed to understand if the early global changes in gene expression were potentiating the later loss of TADs and loops. To do so, we quantified the influence of gene deserts on changes in 3D chromatin structure. If the loss of TADs we observed was less pronounced in gene deserts than actively transcribed genomic regions, it would hint at a direct influence of global changes in transcription on the altered 3D genome organization. First, we defined gene deserts by the absence of active transcription within 20kb regions (Figure 5D). Then, we quantified the insulation score of TADs in gene deserts compared to other TADs. There was no significant difference between these two groups of TADs (Figure 5D). As such, the mechanism behind the loss of TADs and loops at 72 h is likely not driven by global changes in transcription itself. Alternatively, altered expression of a specific set of genes could explain the kinetics and patterns of 3D chromatin changes.

Therefore, we mined our RNA-Seq data to identify key regulators of chromatin structure that were significantly altered from 8 h to 72 h, in a pattern akin to the 3D genome organization changes. One such epigenetic regulator, CTCF, known to promote TAD and loop formation and insulation^42^, displayed a gradual decrease in mRNA and protein levels from 24 h to 72 h, especially accentuated at the later time point (Figure 5E, Figure S6C). Although not gradual, topoisomerase II gene, TOP2A, also displayed a clear downregulation at 72 h (Figure 5F). Similarly to CTCF, the downregulation of TOP2A has also been shown to result in the loss of loops and TADs^43^. Further, using ATAC-Seq, we found that the downregulation of CTCF and TOP2A were also associated with a loss of chromatin accessibility surrounding these genes in GR compared to DMSO condition (Figure S6D). Consequently, CTCF and TOP2A might explain the mechanism of 3D genome organization changes observed in GR treatment.

To test this hypothesis, we investigated whether the changes in chromatin conformation upon CTCF loss or topoisomerase inhibition could explain the marked loss of TAD and loops following GR treatment. Using siCTCF and etoposide, a topoisomerase 2 inhibitor, we knocked down CTCF and inhibited topoisomerase activity in distinct K562 cells (Figure S6E). Then, we performed Hi-C to determine if the changes in loops and TADs were associated with those seen in GR treatment. As expected, APA analysis of the altered loops following siCTCF revealed a marked loss of 2331 loops compared to a few 315 modestly gained loops (Figure 5G). Further, insulation score analysis of TAD boundaries following etoposide treatment revealed a clear loss of insulation at the 1705 lost TADs compared to the control condition (Figure 5H). These results confirm that CTCF downregulation or topoisomerase 2 inhibition led to loss of 3D genome organization.

Next, we aimed to quantify the overlap between the loss of 3D genome organization caused by siCTCF and etoposide compared to GR treatment. Interestingly, a 72% and 65% majority of the 1271 lost TADs and 2332 lost loops in siCTCF conditions were also lost following GR (Figure 5I). Similarly, a 64% majority of the 1705 TADs lost following etoposide treatment were also lost in GR treatment (Figure 5J). When compared together, lost TADs in siCTCF and etoposide treated conditions displayed an additive effect (Figure 5K), which together could, in part, recapitulate and explain the loss of chromatin organization in GR treatment. Therefore, these additional Hi-C experiments highlight that loops and TADs sensitive to the alterations of distinct regulators of 3D chromatin structure also display sensitivity to GR treatment.

Additionally, CTCF and TOP2A downregulation are directly associated with the upregulation of SS target genes caused by GR treatment. Indeed, using the CLUE database from the Broad Institute^44^, we identified genes for which the upregulation displayed the highest connectivity score with the downregulation of CTCF or TOP2A mRNAs. Then, we detected a noticeable overlap between these genes and SS-related genes upregulated by GR treatment and proximal to REST binding sites (Figure 5L-M, Supplementary Text). This analysis supports a mechanism by which GR treatment blocks SS interactions, upregulating their target genes, which in turn promote the downregulation of CTCF and TOP2A. Overtime, this negative feedback loop on epigenetic regulators of 3D chromatin structure plummets in a global loss of TADs and loops, amplifying the upregulation of SS regulated genes and further promoting cell cycle arrest and growth inhibition by GR treatment.

### Combinational treatment of GSK343 and X5050 exerts synergistic antitumor effects

Following our epigenomics analysis of the effect and the mechanism of action of GR treatment, we aimed to quantify the synergy between GSK343 and X5050 in various biological conditions. We calculated the Bliss synergy scores of GSK343 and X5050 in K562^45^, and additionally in the human pediatric B-ALL cell line, SEM^46^. The summarized Bliss scores were 28.809 and 11.382 in K562 and SEM cells, respectively (Figure 6A), which represents a high likelihood of synergy in both models.

**Figure 6.**
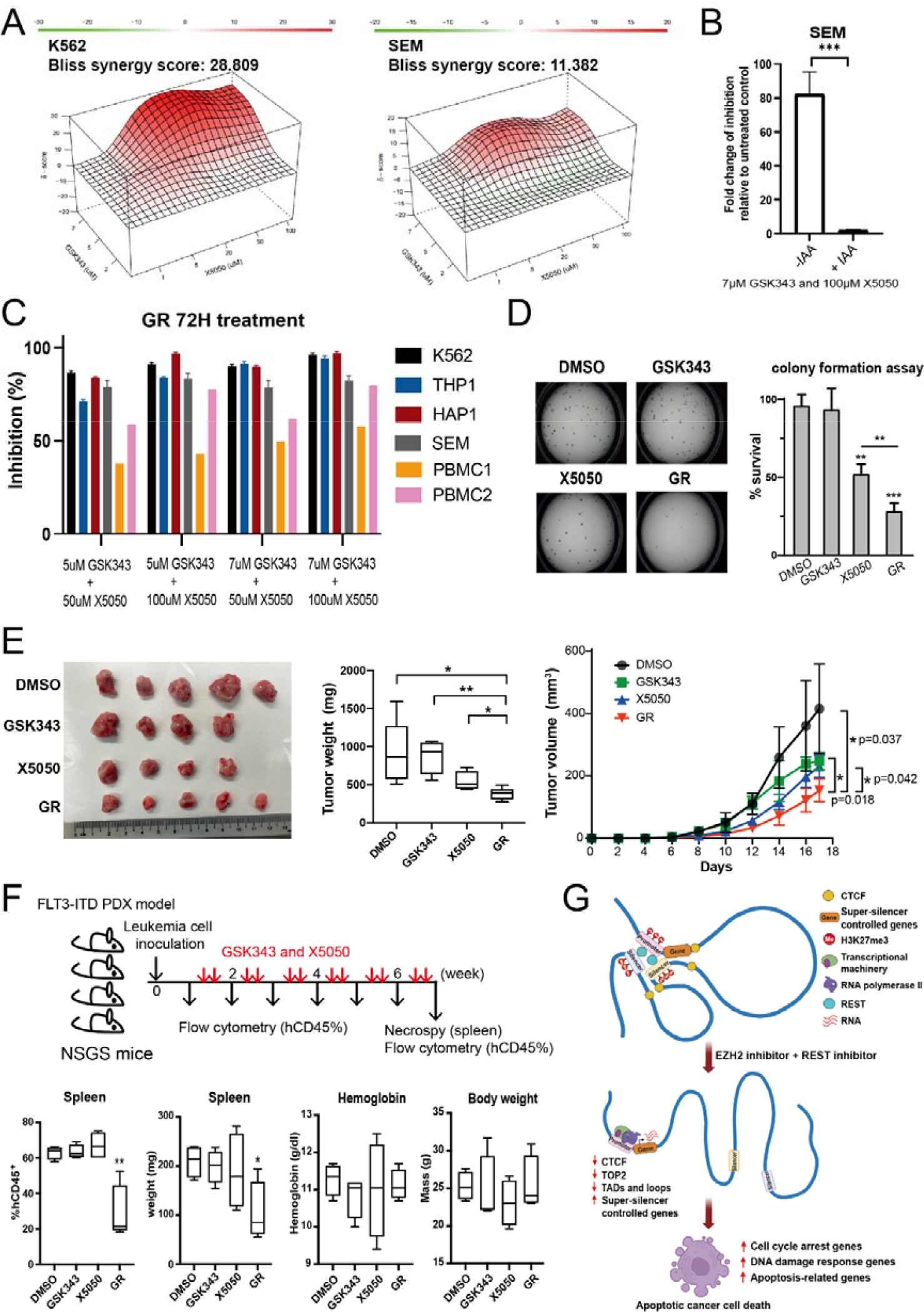
Combinational treatment of GSK343 and X5050 exerts synergistic antitumor effects. **A**. 3D drug interaction landscapes based on Bliss model shown for GSK343 and X5050 combination in K562 cells (left) and SEM cells (right). Summarized synergy scores are calculated across all tested concentration combinations of GSK343 and X5050. Summarized Bliss score above 10 indicates two drugs likely to be synergistic. **B**. Bar chart showing fold change in growth inhibition relative to vehicle control in SEM cells either pre-treated with or without IAA and doxycycline, followed by treatment of GSK343 (5 µM) and X5050 (100 µM) for 72 h. IAA-inducible degradation system is employed to rapidly deplete CTCF protein in SEM cells following 48-h treatment of IAA (50 µM) and doxycycline (1 ug/ml). **C**. Bar chart showing percentage of growth inhibition in four cancer cell lines (K562, THP1, HAP1 and SEM) and two PBMCs. Four different GR conditions are tested for 72 h. **D**. Representative images of the colony formation after being treated with DMSO, GSK343, X5050 or GR for 14 days (left). Bar chart showing percentage of survival in K562 cells treated with DMSO, GSK343, X5050 or GR for 14 days (right). **E**. Representative tumors pictured at final day (left), tumor weight (mg) at final day (middle) and tumor volume (mm^3^) with different post implantation days (right) shown. Tumor growth in NSG mice (NOD scid gamma mice) injected with K562 cells together with different drugs (DMSO, GSK343, X5050 and GR). N=5 for each group. **F**. Boxplots showing percentage of CD45-positive cells in spleen, spleen weight, hemoglobin and body weight at endpoint. GSK343 and X5050 administrated against PDX AML29 cells. 0.1 mg/kg GSK343 and 0.25 mg/kg X5050 injected as either single or combination treatment, twice a week intraperitoneally, after inoculation of leukemia cells (n=4, each group). **G**. Schematic depicting combinational treatment of EZH2 inhibitor (GSK343) and REST inhibitor (X5050) leading to apoptotic cancer cell death. Data shown as average + SEM. P value less than 0.05, 0.01 or 0.001 shown as *, ** or ***, respectively.

Next, as we highlighted the involvement of CTCF in the epigenetic mechanism of action of GR, we investigated the biological influence of its expression on the synergistic effect of the combination treatment. We employed the auxin (IAA)-inducible system to acutely deplete CTCF protein in SEM cells^46,47^. In IAA+ cells, which adapted to the depletion of CTCF (Figure S7A), GR treatment led to a modest but significant 2.1-fold increase in growth inhibition relative to the untreated IAA+ control (Figure 6B). Comparatively, GR treatment in IAA-, where CTCF is not depleted, led to an 82.3-fold increase in growth inhibition compared to untreated IAA-(Figure 6B). Therefore, these results confirm that the depletion of CTCF caused by GR plays an important role in its mechanism of action, while highlighting that cells which are already adapted to low levels of CTCF are relatively resistant to the combinational GR treatment.

To determine if cancer cells were more vulnerable to the combination treatment, we tested different GR concentrations for 72 h in a panel of leukaemia cells, including K562, THP1, HAP1, and SEM, and two peripheral blood mononuclear cells (PBMCs) (Figure 6C). All leukaemia cells tested showed a similar sensitivity to GR treatment as K562 cells, while the two normal PBMCs showed modest inhibition (Figure 6C). This suggested that compared to normal cells, leukaemia cells were more sensitive to GR treatment.

To investigate the synergy of GR is due to a reduction of the REST protein by X5050, we stably knocked down the REST protein in K562 cells and treated these cells with different concentrations of GSK343 (Figure S7B). REST-depleted K562 cells were significantly more sensitive to GSK343 treatment than control cells (Figure S7B), despite the incomplete ablation of REST protein levels. Similarly, we decided to test the synergy between EZH2 and REST inhibition through gene knockout of each gene in distinct HAP1 cells. Then, we treated our HAP1 EZH2 KO cells with X5050 and HAP1 REST KO cells with GSK343 to determine if the combination of genetic KO and single treatment would reconstitute the effect of GR. Indeed, EZH2 KO and REST KO cells were more sensitive to their respective inhibitor treatments, which confirmed that disruption of EZH2 together with REST can lead to synergistic cancer cell death (Figure S7C).

Following our results, which supported the synergy between EZH2 and REST inhibition to repress cancer cell viability, we decided to investigate whether these results would translate *in vivo*. As a preliminary step in determining the antitumoral effect of GR treatment, we tested the effects of single and combination treatments on colony formation assay of K562 cells. Although X5050 treatment alone was able to inhibit colony formation, GR treatment displayed a significantly greater reduction in the number of colonies than all other conditions (Figure 6D). Therefore, the colony formation assay, as for cell viability assay, also supported the synergistic effect of GR, which we proceeded to test *in vivo*. First, we inoculated K562 cells into mice and injected them with control, single, or combination treatments. Consistent with the colony formation assay, single treatment with GSK343 did not show any tumor volume reduction and X5050 significantly reduced the tumor volume. However, here again, GR showed the most potent tumor growth inhibition, being significantly lower than all other conditions (Figure 6E).

Finally, we performed similar treatment conditions in another *in vivo* model with Patient-Derived Xenografts using AML cells carrying a fms-like tyrosine kinase (FLT3)-Internal tandem duplication (ITD) mutation, commonly detected in about 30% of adult AML cases and has bad prognosis^48^. FLT3 inhibition appears to regulate EZH2 to maintain leukemia cells in a stem-like condition^49^. GR treatment was the only condition which displayed significantly reduced cancer burden and hCD45% (Figure 6F). Additionally, we did not detect significant adverse effects in the normal tissues of mice as indicated by the haemoglobin levels and body weight following single or combination treatment (Figure 6F). Together with the observed modest effect on normal PBMC cell viability, our results suggest that the combination of GSK343 and X5050 display a synergistic and cancer-specific antitumoral effect.

In summary, we demonstrated that silencers can work synergistically as “super-silencers”, which are enriched for disease-associated DNA variations (Figure S7D). Further, we showed that super-silencers can be targeted synergistically by inhibiting H3K27me3 deposition by EZH2 and the transcription factor REST. Epigenetically, the combination treatment drives the loss of SS-mediated interactions repressing genes involved in cell cycle arrest and apoptosis, leading to their upregulation and global loss of chromatin structure. In turn, this chain of events leads to synergistic and specific antitumoral effects, which we validated *in vivo* (Figure 6G). Therefore, a combination of epigenetic agents aimed at super-silencers opens promising therapeutic avenues for cancer care.

## Discussion

In this paper, we initially demonstrate that silencers can function synergistically, acting as a super-silencer, to collectively impede gene transcription. Despite our direct demonstration of silencer collaboration for the repression of FGF18, some questions remain unanswered about the mechanism of action of super-silencers across the genome. Depending on the local chromatin structure and transcriptional context, super-silencers may exhibit various forms of synergy. For instance, a single silencing element could be capable of inhibition on its own, while other silencing elements within the SS only potentiates its effect. Therefore, akin to the debate surrounding enhancers and super-enhancers^12,22^, the differentiation between silencers and super-silencers as distinct entities may be subject to discussion. Nevertheless, the demonstrated synergy among silencers is evident and likely to be applicable, albeit with some mechanistic variations, to H3K27me3-rich regions throughout the genome.

We later show that simultaneous inhibition of EZH2, responsible for H3K27me3 deposition, and REST, a known repressive transcription factor, synergise together to promote apoptosis and anti-tumoral effects. Epigenetic silencing of tumor suppressor genes (TSG) is a common hallmark of cancer. This phenomenon has been classically associated to DNA hypermethylation of TSG promoter proximal regions^50^, and has driven the research and implementation of inhibitors of DNA methylation in a wider range of cancer^51^. While an EZH2 inhibitor has been approved for clinical usage^52^, its application is still limited to EZH2 mutated cancers. Our results suggest that EZH2 and REST inhibitors could be utilized more broadly to target oncogenic super-silencer activities in cancers, such as leukemias. Consequently, the therapeutic pathway we propose holds promise for the treatment of a wide range of cancer potentially relying on the unexplored mechanism of super-silencers. Moreover, its combinatorial nature minimizes the likelihood of resistance and reduces the necessity for excessively high, toxic doses to achieve desired effectiveness.

The relationship between 3D genome organization and transcription has been discussed for a long time. 3D genome organization such as TADs are thought to regulate transcription. Transcription may in turn affect the 3D genome organization. Li et al found that, upon blocking RNA polymerase II transcription using flavorpiridol, TAD border strength was reduced, while the inter-TAD interactions increased. Another study in *Drosophila* inhibited transcription using triptolide and showed similar conclusions and more dramatic changes in TADs. Here, we showed that CRISPR knockout of silencers can only change the local TAD structures, but the overall TADs remained similar, although there were genome-wide transcriptional alterations in the KO cells. During the GR treatment experiments, we demonstrated that loss of TADs was not dependent on transcription. Moreover, the early time course GR treatment (8h and 24h) results showed that transcription levels changed early on, but this led to subtle changes of TADs and loops. Our results added on to the evidence that although transcription can affect the 3D genome organization in some contexts, in other contexts, alterations of 3D genome organization are not dependent on the transcription.

Lastly, we showed that the effects of GSK343 and X5050 involved the inhibition of SS interactions, causing the upregulation of SS target genes, which results in major loss of chromatin organization through CTCF and TOP2A downregulation. As the major changes in CTCF, TOP2A and chromatin organization as a whole arise simultaneously with apoptosis, the interplay between this element raises additional questions. CTCF has been shown to display anti-apoptotic effects^53,54^. Similarly, the loss of topoisomerase 2 activity is known to trigger apoptosis, in part due to the genomic accumulation of torsional stress^55^. Additionally, chromatin condense gradually when apoptosis is trigger^56^, but the finer details and kinetics of how chromatin organization is changed during apoptosis is still unknown. Although we outlined specific SS-target genes for which the upregulation is associated with CTCF and TOP2A downregulation, it is possible that currently unknown early apoptotic processes also contribute to the downregulation CTCF and TOP2A to advance apoptosis and the genome-wide loss of 3D genome organization we observed. This hypothesis is further supported by the marked enrichment of SS-target genes upregulated by GR for apoptotic genes in general. As such, our investigation lays the groundwork for further studies of the relationship between epigenetics, chromatin conformation and apoptosis. Future investigations of combination treatments targeting super-silencers, such as GR, could exploit this potential interplay between apoptosis and super-silencers to expand the range of cancers potentially sensitive to this new approach.

In conclusion, our data demonstrated the first example of a SS, constitute of two silencer components that can cooperate to work synergistically via chromatin interactions. Furthermore, we revealed that combinational usage of GSK343 and X5050 could potentially lead to cancer ablation through disruption of SS.

## Methods

We performed Hi-C, ChIP-seq, CUT & RUN, RNA-seq, ATAC-seq, HiChIP, cell culture, RT-qPCR, CRISPR excision, 4C-seq, 3C-PCR, xenograft models, western blot, cell cycle analysis, colony formation assay, adhesion assays, siRNA knock down experiment and growth curves as described in the **Supplementary Methods**. A list of all libraries used and generated is provided in **Supplementary Data 6**. A list of all the primers used is provided in **Supplementary Table S1**.

## Acknowledgements

This research is supported by a National Research Foundation Competitive Research Programme grant awarded to V.T. as lead PI and M.J.F. as co-PI (NRF-CRP17-2017-02). This research is supported by the National Research Foundation Singapore and the Singapore Ministry of Education under its Research Centres of Excellence initiative. This research is supported by a Ministry of Education Tier II grant awarded to M.J.F (T2EP30120-0009). This research is supported by Agency for Science, Technology & Research (A*STAR), National Research Foundation (NRF), Award no. NRF-CRP26-2021-0001 and National Medical Research Council (NMRC), Award No. OFIRG21jun-0101.

## Author contributions

Y.Z., K.J.C., S.C.T., Y.C.C. and M.J.F. conceived of the research and contributed to the study design. Y.C.C. and K.J.C. performed bioinformatics analysis. Y.Z. designed and performed CRISPR knock out experiments, Hi-C for KO, GR treatment and siCTCF, 4C, 3C-PCR, RNA-seq, ChIP-seq, ChIP-qPCR and other functional experiments for KO clones. S.C.T. performed ATAC-seq, Cut & Run assay, cell viability assay and drug combination assay. A.N. and M.O. performed the drug synergy, colony formation assay, cell cycle analysis and mice experiments for the drug treatment. M.M. assisted with RNA-Seq. A.R., L.M. and V.T. designed and performed the xenograft experiments for DKO. Y.X.S. helped with the GSK343 treated Hi-C preparation. C.Y.F. helped with the microscope imaging and analysis. Z.X.L. performed Hi-C experiments for the drug treatment. B.J. performed the connectivity analysis. Y.Z., K.J.C., S.C.T., Y.C.C., B.L., and M.J.F. reviewed the data and wrote the manuscript. All authors reviewed and approved the manuscript.

## Data deposition

The list of libraries used in the study is provided in **Supplementary Data 6**. All datasets have been deposited into GEO under accession number GSE193489. Private access token **apepeyskxbedrsz** can be used to view the private data.

## Author information

The authors declare that they have no competing interests. Correspondence and requests for materials should be addressed to.

## Notes

### Competing Interest Statement

The authors have declared no competing interest.

