## Supplementary text, Supplementary Tables and Supplementary Figures for "Super-silencer perturbation by EZH2 and REST inhibition leads to large loss of chromatin interactions and reduction in cancer growth"

**Supplementary Materials**

### Contents

1. **Supplementary Text**
2. **Methods**
3. **Supplementary Tables**
4. **Supplementary Data**
5. **Supplementary Figures and Legends**
6. **Supplementary References**

### Supplementary Text

#### Integrative analysis of 4C and ChIP-Seq

To dissect the synergistic mechanism systematically and in depth, integrative analysis of 4C-seq and ChIP-seq was performed. Loops were classified into three different categories including gained loops, lost loops, and unchanged loops, and correlated them with histone modifications. There were increased H3K27ac signals and decreased H3K27me3 signals for unchanged loops in DKO compared with EV (Figure S3C-D). Moreover, H3K27ac signals were also increased at gained loops (Figure S3C-D), suggesting the loops being more active in DKO cells. Gained H3K27ac signals and depleted H3K27me3 signals were observed again when comparing DKO to S1KO (Figure S3E-F), but not when comparing S1KO to EV (Figure S3G-H), suggesting differences indeed exist between S1KO and DKO. S1KO did not display many epigenomic differences, possibly due to compensation by S2.

#### Identification of GR upregulated SS genes associated with reduction of CTCF/TOP2

To find the potential GR controlled and SS controlled synergistic upregulation genes that are related to CTCF and TOP2 mRNA reduction, we first shortlisted genes upregulated in GR-treated cells at 72 h. Next, we overlapped these 1952 genes with the potentially SS controlled genes identified by the H3K27me3 HiChIP, which revealed 261 upregulated SS genes (Figure 5L). Among these genes, 193 were associated with REST binding. *FGF18* was one of the genes in this category. Next, we obtained the list of genes that potentially lead to reduction of CTCF and TOP2 expression from Connectivity Map[1]. After overlapping, we found that 19 genes out of 193 were related to CTCF reduction while 40 genes out of 193 were related to TOP2 reduction (Figure 5L). Finally, we shortlisted eight genes that exhibited synergistic upregulation (Figure 5M). The fold changes in gene expression for these genes were greater in GR-treated samples compared to the sum of GSK343-treated and X5050-treated samples. Furthermore, these upregulated genes were linked to reduced levels of CTCF and TOP2 mRNA, as well as the loss of TADs at 72 hr (DMSO vs GR).

### Methods

#### Cell culture

Human chronic myelogenous leukemia cell line K562 and human leukemia monocytic cell line THP1 were cultured in RPMI-1640 supplemented with 10% Fetal Bovine Serum (FBS) and 1% penicillin-streptomycin. HAP1, a near-haploid human leukemic cancer cell line, was cultured in IMDM supplemented with 10% FBS and 1% penicillin-streptomycin. SEM CTCFAID cell line was a kind gift from the lab of Dr. Li Chunliang[2]. SEM cells were cultured in RPMI 1640, 2 mM L-glutamine, 10% FBS, and

1% penicillin-streptomycin. All cultures were maintained at $37^{o}C$, 5% CO2 in a humidified incubator.

**CRISPR excision**

CRISPR excision was performed using the all-in-one CRISPR/Cas9 vector system, as described previously[3]. Briefly, gRNAs were designed using Zhang Feng's website (http:// CRISPR.mit.edu)[4]. Two gRNAs were designed to perform the excision for each targeted region. Single gRNA was cloned into either pX330A/pX330S vector (gift from Li Shang, pX330A modified to include GFP reporter marker) followed by Sanger sequencing to confirm correct insertion of gRNA. Golden gate assembly of two gRNAs was performed as previously described to put them into the pX330A backbone. Two positive gRNAs insertion plasmids were confirmed by Sanger sequencing using CRISPR-step2-F and CRISPR-step2-R primers (sequences shown in Table S1). Next, the plasmid was electroporated into the K562 cell line using the Neon transfection system (Thermo Fisher). After 48 h, transfected cells were FACS sorted into 96-well plates as single-cell colonies based on GFP signal.

Cells were harvested from each clone, pelleted and lysed in lysis buffer. Genotyping was performed using an internal and flanking primer pair. Final PCR products were imaged by agarose gel electrophoresis and successful clones were confirmed through Sanger sequencing (First Base). All primers used for genotyping were listed in Table S1.

**RNA extraction and RT-qPCR**

Total RNA was isolated from cells using RNeasy Mini Kit (Qiagen) and on-column DNase digestion (Qiagen) was performed. RNA concentrations were determined by Nanodrop ND1000 (Thermo Scientific) and 1 µg of total RNA were then reverse transcribed to cDNA using the SuperScript III first-strand synthesis system (Invitrogen). Quantitative real-time PCR (qPCR) was then performed using the Applied Biosystems QuantStudio 3/5 Real-Time PCR system using SYBR Green PCR Master Mix and appropriate primers listed in Table S1. Gene transcript levels were analysed by $2^{-\Delta\Delta Ct}$ method[5].

**RNA-seq**

The total RNA was isolated as described in RNA extraction session. Quality of the extracted RNA was analysed using Agilent RNA 6000 Nano Kit (Agilent) and quantitated using Nanodrop ND1000 (Thermo Scientific). Library construction was performed using TruSeq Stranded Total RNA LT (with Ribo-Zero Gold) Set A (Illumina), as per protocol. Libraries were then sequenced using the Illumina HiSeq4000 platform.

**Adhesion assay**

Cell adhesion assay was performed using CytoSelect 48-Well Adhesion Assay (Cell Biolabs, San Diego, CA), according to the manufacturer's protocol. Briefly, a cell suspension ($5X10^5$ cells in 200 ml FBS-free medium) was added to fibronectin-coated wells and BSA coated wells (negative control), respectively. After 3 h of incubation, cells were washed with PBS, stained with crystal violet and eluted with extraction solution. Adhesion levels were quantified by optical absorbance at 560 nm using the Tecan plate reader.

**Growth curve assay**

1000 cells/well were seeded in 96 well plates. Cell growth was measured at day 0, day 1, day 2, day 4 and day 5 by CellTiterGlo assay kit (Promega, G7571). Luminescence was read on a Tecan plate reader.

**Xenograft experiments**

All the animal studies were carried out in accordance with animal care and use guidelines approved by Biological Resource Centre, Singapore.

Six to eight weeks old female CB17 SCID mice were used for the present study. The mice (n=5) were injected subcutaneously. All the mice were monitored for tumor growth at the inoculation site and tumor volume was measured using Vernier caliper twice weekly for 40 days or until tumor volume reached 1000 $mm^3$, whichever was earlier. Tumor volume was calculated using the formula $V = a \times b^2 \times 0.52$, where a represents the largest diameter and b represents the smallest diameter of the tumor.

The xenograft model presented in Figure 1F were generated using the following method: one of the DKO CRISPR clones (“DKO-C1”) and its empty vector clone (“EV”) were injected in the same mice (left side and right side), while another DKO CRISPR clone (“DKO-C2”) and its EV were injected in the same mice (left side and right side) to reduce the tumor growth differences in different mice.

The xenograft model presented in Figure 6E was generated using the following method: NSG mice (NOD scid gamma mice) were inoculated subcutaneously in the left flank with 5 $x10^6$ K562 cells, and randomized into four groups: vehicle, GSK343, X5050, and GR. In groups treated either a single agent or combination treatment, the same doses of individual drugs (0.2 mg/kg GSK343 and 0.5 mg/kg X5050) were administered via intraperitoneal injection twice weekly. Tumor sizes were serially measured by digital caliper and tumor volumes were estimated according to the equation $V = 1/2$ (length x width$^2$). At the endpoint, individual tumors were excised, weighed and recorded by photograph.

The serially transplantable PDX model presented in Figure 6F was generated using the following method: $4x10^6$ AML29 PDX human leukemia cells carrying FLT3-ITD mutation were transplanted into 2 Gy irradiated NSGS mice by intravenous injection and randomized into four groups (vehicle, GSK343, X5050, and GR). In groups treated either a single agent or combination treatment, the same doses of individual drugs (0.1 mg/kg GSK343 and 0.25 mg/kg X5050) were administered via intraperitoneal injection twice weekly, starting from 2 days after transplantation and continuing for 7 weeks. At the endpoint, spleen was excised and weighed. Cells in bone marrow (BM), spleen and peripheral blood (PB) were harvested and analyzed by flow cytometry.

**siRNA knockdown experiment**

siRNAs (Thermo Fisher) were electroporated into DKO cells using Neo transfection system (Thermo Fisher) according to the manufacturer’s instructions. *FGF18*-siRNA-2, 5’-GAGACGGAAUUCUACCUGUtt-3’, *FGF18*-siRNA-3, 5’-AGACACCUUCGGUAGUCAAtt-3’, were used for siRNAs targeting *FGF18* genes. After 48 h incubation, cells were harvested for RT-qPCR and seeding for growth curve measurement. CTCF siRNA knockdown was performed using ON-TARGETplus SMARTpool Human CTCF (10664) siRNA (Dharmacon L-020165-00-0005). After 48 h incubation, cells were harvested for western blot and Hi-C. Primers used for RT-qPCR were listed in Table S1.

**shRNA knock down experiment**

Two shRNAs against REST gene (shREST-3: 5’-CCGGGCATCCTACTTGTCCTAATACTCGAGTATTAGGACAAGTAGGATGCTTTTTG-3’ and shREST-4: 5’-CCGGGCCTCTAATCAACATGAAGTACTCGAGTACTTCATGTTGATTAGAGGCTTTTTG-3’) were cloned into pLKO.1-TRC vector (Addgene: 10878). Plasmids were transfected into K562 cells by lentiviral transduction. Stable REST knockdown K562 cells were selected with puromycin for three days.

**Circular chromosome conformation capture sequencing (4C-seq)**

4C-seq was performed as previously described but with slight modifications[6]. Briefly, 40 million cells were cross-linked with 1% formaldehyde. Nuclei pellets were isolated by cold lysis buffer (10 mM Tris-HCl, 10 mM NaCl, 5 mM EDTA, 0.5% NP-40) supplemented with protease inhibitors (Roche). First digestion was performed overnight at 37°C with HindIII enzyme (NEB). Digestion efficiency was measured by gel electrophoresis. After confirmation of good digestion efficiency, DNA was ligated overnight at 16°C by T4 DNA ligase (Thermo Scientific) and de-crosslinked by proteinase K. Then, DNA was extracted by phenol-chloroform, referred to as “3C library”. “3C library” was then processed overnight at 37°C for second digestion with DpnII enzyme (NEB). After ligation, “4C template DNA” was obtained. Concentration of 4C DNA was determined by Qubit assays (Thermo Scientific). 4C template DNA was then amplified using specific primers with Illumina Nextera adapters and sent for sequencing on the MiSeq platform. Primers used for 4C amplification were listed in Table S1.

**3C-PCR**

The “3C library” was generated as described in the 4C-seq section. After confirmation of good digestion efficiency by HindIII, Taq PCR core kit (Qiagen) was used for PCR reactions. Briefly, 600-800 ng “3C library” was measured by Qubit assays (Thermo Scientific) and used the following PCR protocol with specific 3C-PCR primers: 98°C 3 min, 33 cycles [94°C 1 min, 60°C 1 min, 72°C 20 sec], and 72°C 10 min. PCR products were run on 1.5% agarose gels. Primers were designed for 3C-PCR following the unidirectional strategy[7]. After gel electrophoresis, bands corresponding to expected products were gel excised and purified (Qiagen) for Sanger sequencing. Band intensities were measured by Image Lab. Primers used were listed in Table S1. At least two replicates were performed for 3C analyses.

**Hi-C experiment and library preparation**

One million cells were fixed and used for Hi-C experiment using Arima-Hi-C kit (Arima Genomics) according to manufacturer’s protocol. Libraries were prepared using KAPA Hyper-Prep Kit (KAPA), according to Arima-Hi-C kit protocol. Final Hi-C libraries were sequenced pair-end 2X150bp on the Illumina HiSeq 4000 platform.

To map chromatin interactions induced by the apoptosis-inducing drug etoposide, K562 cells were either treated with 0.1% DMSO or 10 μM etoposide for 72 h and processed for Hi-C library preparation.

**ChIP-seq and ChIP-qPCR**

ChIP-seq was performed according to Robertson *et al.* with slight modifications[8]. Briefly, cells were fixed with 1% formaldehyde (Thermo Scientific) at room temperature for 10 min, followed by quenching with glycine for 5 min at room temperature. Fixed cell pellets were lysed in 1% SDS lysis buffer supplemented with protease inhibitor cocktail tablet (Roche) and sonicated using Bioruptor (Diagenode).

Cell lysate was precleared through centrifuging in dilution buffer and incubating with Protein G Dynabeads (Invitrogen) overnight at 4°C. Precleared lysate was added into the antibody-conjugated beads and incubated overnight at 4°C, with rotation. The incubated beads were then washed thrice by 0.1% SDS lysis buffer, once with high salt wash buffer, once with lithium chloride and once with TE buffer. The incubated beads were then added into the elution buffer with RNase A (Qiagen), followed by decrosslinking with Proteinase K (Ambion) overnight at 37°C. ChIP DNA was extracted with the QIAquick PCR purification kit (Qiagen) and quantitated using Qubit High Sensitivity dsDNA Assay (Invitrogen).

ChIP-seq library construction was prepared using ThruPLEX DNA-seq 48D Kit (Rubicon), in accordance with instructions and the prepared library was then sequenced at the Illumina HiSeq 4000 platform. Antibodies used include H3K27me3 (C36B11, Cell Signaling Technologies), H3K27ac (#ab4729, Abcam) and mouse IgG (#sc-2025, Santa Cruz). 3.5 µg of antibodies were used for each ChIP.

ChIP-qPCR reactions were performed with the QuantStudio 5 quantitative PCR machine (Life Technologies) using GoTaq qPCR Master Mix (Promega) by at least three ChIP DNA replicates. Primers used were listed in Table S1.

#### Protein extraction and western blot

Cells were lysed using RIPA buffer (Sigma-Aldrich) with protease inhibitor cocktail (Life Technologies) at 4°C for 30 min. Cell lysate was centrifuged to remove cell debris and protein concentration was determined by Pierce BCA Protein Assay Kit (Thermo Scientific). 20 µg of proteins were separated in 4-20% Mini-PROTEAN® TGX™ Precast Gels (Bio-Rad) and transferred to PVDF membrane. After blocking with TBST containing 5% nonfat dried milk at room temperature for 1 h, membrane was washed with TBST and incubated overnight with primary antibodies at 4°C: EZH2 (Cell Signaling Technology 3147), beta-Actin (abcam ab6276), REST (Merck Millipore 17-641, Proteintech 22242-1-AP), cleaved-PARP (Cell Signaling Technology 9541S), gamma-H2AX (Merck Millipore JBW301), and CTCF (Santa Cruz sc-271474). Membrane was washed three times with TBST for 10 min each time and then incubated at room temperature for 1 h with HRP-conjugated secondary antibodies (Cell Signaling Technology). After extensive washing, bands were detected by chemiluminescence reagent (Bio-Rad) and imaged using ChemiDoc™ imaging system (Bio-Rad). Protein levels were calculated by Image Lab.

#### Drug combination assay

K562 and SEM cells were seeded on 96-well culture plates at a density of 4000 cells/well and treated with GSK343 (0, 2, 5, and 7 µM) and X5050 (0, 1, 5, 20, 50, and 100 µM) for 72h as either a single agent or in combination. To rapidly deplete CTCF in SEM cells, cells were seeded as described above[2], and pre-treated with IAA (50 µM) and doxycycline (1 µg/ml) for 48 h prior to the addition of GSK343 and/or X5050. Following 72 h incubation, cell viability was measured by CellTiter-Glo assay kit (Promega, G7571) after 72 h incubation. Luminescence was read on a Tecan plate reader. Dose-response matrix of GSK343 and X5050 in K562 and SEM cells was shown by % inhibition. The Bliss synergy score for GSK343 and X5050 was calculated using SynergyFinder 3.0 (https://synergyfinder.fimm.fi/)[9]. A summarized Bliss synergy score above 10 indicates that two drugs are likely to be synergistic.

**Cell cycle analysis**

Cell cycle analysis was performed 24 h after drug treatments with GSK343 and/or X5050 using BrdU flow kit (BD Pharmigen, NJ) according to manufacturer's instructions. Briefly, K562 cells were labelled by BrdU for 2 h before fixation and incubated for 30 min at room temperature with 200 ng/ml RNaseA. Next, DNA was stained with 25 ug/ml PI and FITC-conjugated anti-BrdU antibody and subjected to flowcytometry LSRII (BD Biosciences, CA).

**Annexin V/PI apoptosis assay**

Annexin V/PI apoptosis assay was performed on DMSO-, GSK343-, X5050- and GR-treated K562 cells according to manufacturer's protocol (FITC Annexin V apoptosis detection kit with PI, BioLegend USA)[10]. Briefly, the K562 cells were washed twice with cold BioLegend's cell staining buffer and then resuspended in Annexin V Binding Buffer at a concentration of 0.5 x $10^6$ cells/ml. Cells were then mixed and 100 µl of cell suspension was passed through a cell strainer, followed by 5 µl of FITC Annexin V and 10 µl of PI staining solution being added to the tube. The cells were gently vortexed and incubated in the dark for 15 min at room temperature. 400 µl of Annexin V Binding buffer was then added to each tube, followed by the samples being analyzed by BD™ LSRII flow cytometer (BD Biosciences, USA). Each sample was tested in triplicates.

**Colony formation assay**

K562 cells were plated in triplicate at a density of 200 cells per dish in 35-mm dishes containing methylcellulose semisolid medium Methocult H4034 (StemCell Tec., Vancouver, Canada). The cells were treated with DMSO, GSK343, and/or X5050. After 14 days of incubation, colonies were counted using an SZX12 Olympus microscopy (Olympus, Japan).

**ATAC-seq**

ATAC-seq was performed using the Diagenode ATAC-seq Kit (Diagenode) following the manufacturer's protocol[11]. Briefly, 200,000 cells of EV, S1KO, S2KO, and DKO K562 cells per sample were harvested and nuclei were isolated. Immediately following the nuclei prep, the pellet was resuspended in 50 µl tagmentation mix (25 µl 2x Tagmentation Buffer, 2.5 µl tagmentase, 0.01% digitonin, 0.1% Tween 20, 16.5 µl PBS and 5.25 µl nuclease-free water). The transposition reaction was carried out for 30 min at 37°C. The tagmented DNA was then purified and eluted in 12 µl of DNA Elution Buffer. Following purification, we amplified library fragments using 10 µl of tagmented DNA and 1 µl of primer pair provided by the 24 UDI for Tagmented libraries Set I (Diagenode) using the following PCR conditions: 72°C for 5 min, 98°C for 30 s, and thermocycling at 98°C for 10 s, 63°C for 30 s and 72°C for 1 min. We first amplified the libraries for five cycles, after which we took 5 µl of amplified DNA and added 10 µl of qPCR mix with Sybr Green. We ran this reaction for 20 cycles to determine the additional number of cycles needed for the remaining 45 µl reaction. The libraries were amplified for additional 6-11 cycles and cleaned up using AMPure XP beads (Beckman Coulter). The indexed libraries were sequenced paired-end 2 x 50 bp on HiSeq X Ten platform.

The same ATAC-seq protocol was used for K562 cells either treated with 0.1% DMSO or GR for 72h with the following modifications. The number of viable cells was manually counted using a hemocytometer and 50,000 nuclei were prepared. The tagmented libraries were amplified for additional 6-7 cycles on the thermocycler and validated using the Agilent High Sensitivity DNA kit. The final libraries were sequenced paired-end 2 x 50 bp on NovaSeq 6000 platform.

**Cut & Run assay**

Cut & Run was performed for CTCF and H3K27me3 using the EpiCypher CUTANA Cut & Run kit according to the manufacturer's instructions (EpiCypher)[12,13]. All the Cut & Run experiments were performed using 500,000 cells, 0.5 µg of antibody, and 10 µl of activated concanavalin A beads per sample. For Cut and Run CTCF, EV, S1KO, S2KO, and DKO K562 cells were harvested and bound to activated concanavalin A beads. Cells were subsequently permeabilized with 0.01% digitonin and incubated with CTCF antibodies (EPR18253, Abcam) at 4°C overnight.

For Cut & Run H3K27me3, K562 cells were either treated with 0.1% DMSO or GR for 24 h. The number of viable cells was manually counted using a hemacytometer and 500,000 cells per sample were harvested. Following digitonin permeabilization, cells were incubated with H3K27me3 antibodies (C36B11, Cell Signaling Technology) at 4°C overnight. The K-MetStat Panel was spiked into each sample as an internal control for H3K27me3 antibody specificity. All libraries were amplified using 14 cycles on the thermocycler and validated using the Agilent High Sensitivity DNA kit. The final libraries were size selected using AMPure XP beads (Beckman Coulter) twice to remove DNA fragments <150 bp. The indexed libraries for CTCF and H3K27me3 were sequenced pair-end 2 x 150 bp on Illumina NovaSeq 6000 platform.

**Human recombinant FGF18 protein assay**

K562 cells were counted and seeded (4000 cells per well) in 96-well plates. Cells were serum-starved for 24 h before addition of rhFGF18. Cells were treated either with sterile water or increasing concentrations of rhFGF18 (0, 10, 50, 100, and 200 ng/ml). After 72 h of treatment, viable cells were measured with CellTiter-Glo assay kit (Promega, G7571). Luminescence was read on a Tecan plate reader.

**RNA-seq, ChIP-seq & 4C data analysis**

The sequencing reads of the RNA-seq libraries for EV, S1KO, S2KO and DKO were quantified using kallisto (0.45.1). The differentially expressed gene analysis was performed using sleuth (0.30.0)[14].

Adaptors were trimmed by trimmomatic (0.39)[15]. After adaptor trimming, reads were mapped to hg19 and processed using STAR (2.7.3a)[16]. Reads normalization and differential expression analysis were performed using DESeq2[17]. To filter out genes, we selected those that were upregulated and downregulated genes with a Log2(fold change) between -1< and 1 and a P value less than 0.05. This filtering method was applied to the RNA-seq results of drug treatments (GSK343, X5050 and GR) depicted in Figure 4-5.

Gene Set Enrichment Analysis (GSEA) was performed using the GSEA pipeline from the Broad Institute[18]. ChIP-seq adaptors were trimmed by trimmomatic (0.39)[15]. The ChIP-seq reads of H3K27me3 and H3K27ac were then mapped by BOWTIE2 (v2.2.5) using default parameters in pair-end mode and filtered out alignment with a mapq score smaller than 30[19]. Two replicates of ChIP-seq were combined, and peaks and bigWig files were generated by MACS2 (2.1.0.20150731) using option "-q 0.01" for H3K27ac, and "–broad –broad-cutoff 0.1 -q 0.05" for H3K27me3[20].

4C reads were trimmed at HindIII digestion site using tagdust (2.33)[21], and only the read pairs that remained after trimming were mapped using BOWTIE2 (v2.2.5) in the single-end mode with option "–end-to-end"[19]. Significant interactions were called against the Hind III digested genome background using R3Cseq (1.24.0) with a cut-off p value of 0.05[22]. Two replicates were performed for each 4C analysis, and significant interactions from the two replicates were pooled. 4C interactions were drawn in arc style using Sushi (1.16.0) from Bioconductor[23].

### Hi-C data analysis

The paired-end reads were mapped to hg19 reference genome using BWA mem (0.7.17-r1188)[24]. The mapped reads were then passed to Juicer pipeline (1.5.7) to perform chimeric reads analysis, merging, generating Hi-C matrix, loop calling using HiCCUPS, and TAD calling using Arrowhead[25]. Insulation scores were calculated by hicFindTADs from HiCExplorer (3.4.2/3.6) using a 10kb resolution matrix[26,27]. The eigenvector values at 1Mb and 50kb resolution, and aggregate peak analysis (APA) at 5kb resolution were performed using the eigenvector and APA function of Juicer tools[25].

For drug treatment K562 Hi-C, the TAD and loops integrative analyses were performed. We divided the TADs and loops into three categories (gained, lost, and unchanged) according to the following principles (use GSK343 treated Hi-C as an example): 1) gained TADs were those TADs only found in GSK343-treated Hi-C libraries; 2) lost TADs were those TADs only found in DMSO-treated Hi-C libraries; 3) unchanged TADs were those TADs found in both GSK343-treated and DMSO-treated Hi-C libraries. To get the classification of gained, lost, and unchanged TADs and loops, we also defined the criteria for determining whether two TADs or loops were the same. If two TADs from different samples reciprocally overlap ratio are more than 50%, we considered as the same TAD. If not, they were considered as different TADs. For loops, if two loops from different samples have at least one base pair overlap, we considered them as the same one. If not, they were considered as two different loops.

APA plots were generated by the APA function in Juicer tools[25]. Briefly, 50kb windows on loops in different categories were visualized using R to make the results of different samples under the same color range. The mean insulation scores of gained, lost or unchanged TADs in each comparison group in K562 drug treatment Hi-C were calculated and visualized to compare the different categories after drug treatment. The mean insulation scores of TADs in K562 FGF18 silencers knockouts (S1KO, S2KO and DKO) were compared using TADs from EV as the reference. All calculation and visualization of insulation scores were done by computeMatrix reference-point and plotProfile from deeptools (3.5.1)[28].

The eigenvector values at 1Mb and 50kb resolution were performed using the eigenvector function in Juicer tools[24]. Density plots of eigenvector value 1 at 1Mb resolution in the whole genome for each comparison group were drawn to show the correlation between different drug treatment Hi-C to DMSO Hi-C. The R in figures obtained from cor.test function in R shows the correlation coefficient of two samples. A correlation coefficient closer to one indicates a higher similarity in the compartments of two samples. An example of chr1 eigenvector values in K562 drug treatment Hi-C at 50kb resolution was shown.

### Heatmap and boxplot of RPM signal and ChIP-seq signal at different 4C regions

4C regions were classified as gained, lost, or unchanged based on their presence in experimental condition versus control condition. Here, we compared three different conditions (DKO vs EV, S1KO vs EV and DKO vs S1KO). The classification criteria were as follows: Gained - 4C interactions only present in the experiment condition but not in the control condition; Lost - 4C interactions only present in the control condition but not in experiment condition; Unchanged - 4C interactions present in both control and experiment conditions. Only 4C interactions with p value < 0.05 were considered in the comparison.

For RPM signal heatmap, different types of 4C regions (gained/lost/unchanged) were first classified based on their H3K27ac/H3K27me3 ChIP-seq signal levels in the control condition. Tertiles were used to assign these 4C regions into different categories (high, medium, and low). The 4C intensity in RPM of each 4C regions were shown in a color-scaled manner.

For the ChIP-seq signal heatmap, the ChIP-seq signals of different types of 4C regions (gained/lost/unchanged) were calculated as the area of signal at these regions. Deeptools computeMatrix was used to calculated ChIP-seq signal at these 4C regions[29], and then the total signal areas were calculated using the formula: Total Signal Area = sum (Sig *BS), where BS represented the size of bins used when summarizing RPKM, and Sig represented the ChIP-seq signal in RPKM at individual bin. For ChIP-seq signal boxplot, the same 4C region (gained/lost/unchanged) in different conditions were connected using gray lines. Wilcoxon paired test was used, and p values were indicated accordingly: ns, p > 0.05, *p < = 0.05, **p < = 0.01, ***p < = 0.001, ****p < = 0.0001.

### K562 H3K27me3 HiChIP and data analysis

HiChIP was performed using H3K27me3 antibody (C36B11, Cell Signaling Technologies) in normal K562 cells. Five million cells were fixed and used for HiChIP experiment using Dovetail HiChIP MNase kit (Dovetail Genomics) according to the manufacturer's protocol. Briefly, 1 µl MNase enzyme mix and 1250 ng H3K27me3 antibody were used for HiChIP experiment. Libraries were prepared using the Dovetail Primer Set for Illumina and Dovetail Library Module for Illumina kits, according to the manufacturer's protocol. The final HiChIP libraries were sequenced pair-end 2X150bp on the Illumina HiSeq 4000 platform.

HiChIP analysis was done by following the documentation of Dovetail website (https://hichip.readthedocs.io/en/latest/index.html) step by step. Paired-end reads were aligned to hg19 using BWA mem (0.7.17-r1188) with parameter "-5SP -T0" to get mapped reads in the SAM file format[24]. Next, the mapped reads were processed by pairtools (0.3.0) (https://github.com/mirnylab/pairtools) to perform valid ligation recording, sorting, removing PCR duplicates, and splitting. After this step, pairs and bam were obtained from pairtools. Bam file was then sorted and produced index file using SAMtools for downstream analysis[30]. Before the further process, the quality of HiChIP library and the enrichment of HiChIP reads at H3K27me3 binding sites were evaluated by the QC report generated from get_qc.py and enrichment_stats.sh scripts in HiChIP (https://github.com/dovetail-genomics/HiChiP/blob/main/docs/source/before_you_begin.rst).

After Passing the QC assessment, pairs file from pairtools was converted into contact matrix in hic format by Juicer Tools and visualized in Juicebox (2.1.10)[31]. HiChIP loops were called by FitHiChIP (9.0) in 25kb bin size[32], and the coverage of the whole genome was calculated by bamCoverage from Deeptools (3.5.1)[31]. Then results at certain regions were visualized using sushi in R[23].

### ATAC-seq analysis

Adaptors were trimmed by TrimGalore (0.6.6) (https://www.bioinformatics.babraham.ac.uk/projects/trim_galore/) and then aligned in pair-end mode using bowtie2 (v2.3.5.1) 13 with default parameters. The resulting SAM files were converted to BAM format and filtered to exclude reads with a mapq score below 30 using samtools (v1.10) 24. The BAM files were sorted, and duplicate reads were removed using through Picard (https://github.com/broadinstitute/picard, v 2.27.5-4). After removing reads mapped to chrM, chrUn, and 'random' chromosomes, BAM files from replicates were merged, then converted to BigWig files with RPKM normalization using deeptools bamCoverage (v3.4.3) 22 for data visualization on the UCSC Genome Browser. Peaks were called using MACS2 (v2.2.7.1) (PMID: 24743991) callpeak with the following flag parameters: --shift -100 --extsize 200 --nomodel -B --SPMR -g hs, and peaks contained in the ENCODE (v2) hg19 blacklist were removed.

**Cut & Run analysis**

The paired-end reads were aligned with bowtie2 (v 2.3.5.1) using the following flag parameters: --local, --very-sensitive-local, --no-unal, --no-mixed, --no-discordant, --phred33, -I 10, and -X 700[18]. The resulting SAM files were then converted to BAM files and sorted with the samtools (v 1.10) and Picard (https://github.com/broadinstitute/picard, v 2.27.5-4) software[29]. After removing reads mapped to chrM, chrUn, and 'random' chromosome, BAM files of replicates were sorted and combined using the samtools (v 1.10) sort and merge function[29]. Merged BAM files were converted to BigWig files with RPKM normalization using deeptools bamCoverage (v 3.4.3) for data visualization on UCSC Genome Browser[27]. Peaks were called using MACS2 (v 2.2.7.1) (PMID: 24743991) with default parameters, and only peaks not contained in the ENCODE (v2) hg19 blacklist were retained.

**Disease-associated SNP analysis**

Three different categories of SNPs lists, namely disease-associated, cancer-associated, and AML/CML-associated SNPs, were downloaded from GWAS[33]. Then the overlapped ratio was calculated in both MRR and SNPs. To confirm the enrichment of SNPs in MRRs, random data with the same number and size of regions as MRR peaks from the genome were created 1000 times. Mean overlapped ratios between random data and SNPs were calculated and the Wilcoxon test was used to compare the difference between random and MRR ratios.

**ChIP-seq peaks of REST at different categories of TADs and loops**

We calculated the number of REST ChIP-seq peaks (https://www.encodeproject.org/experiments/ENCSR137ZMQ/) at gained/lost/unchanged TADs region in GSK343 vs DMSO. ChIP-seq peaks with at least one overlapped base pair with the TADs were counted and used to generate the violin plot by R (Figure 3E).

**Gene deserts analysis**

Intergenic regions were identified based on known gene locations. If the size of intergenic regions ⩾ 500kb, these regions were defined as gene deserts. TADs that overlapped with gene deserts were considered as "TADs in gene deserts" and compared with all the TADs in the following manner: We set the start and end positions of TADs extended with 5kb upstream and downstream as the TAD boundaries. Then we calculated the mean insulation score differences of the TAD boundaries for two categories (all TADs and TADs in gene deserts) in the comparison between GR vs DMSO at 72 h. A box plot was generated to visualize the change in insulation scores. Wilcoxon test was used to evaluate the difference between all TADs and TADs in gene deserts.

The RNA-seq reads in DMSO and GR at 72 h were calculated in each 20kb bin across the whole genome and gene deserts. These reads were then normalized by the total number of mapped reads in the whole genome as RNA signals. Boxplots were generated to compare the normalized RNA-seq reads between whole genome and gene deserts. Wilcoxon test was used to compare the difference between them.

### Supplementary Tables

**Table S1.** Primers used for ChIP-qPCR, RT-qPCR, 4C-Seq, 3C-PCR and CRISPR genotyping primers and guide RNAs used for CRISPR.

| Primer name | Sequence (5' to 3') |
|---|---|
| ChIP-qPCR primers | |
| FGF18-R1-ChIP-F | AACCTGCTTTGAGTTTTCGGC |
| FGF18-R1-ChIP-R | CAAGATGCCGGGAACAGCAC |
| FGF18-R2-ChIP-F | AACAGAGCTCCCGATGAACC |
| FGF18-R2-ChIP-R | GAGACGCTTCCCTGTCACTG |
| FGF18-R3-ChIP-F | TTCGGGGTAGGACGTGTCA |
| FGF18-R3-ChIP-R | GGCTCGGAAACAAGGTCTGT |
| FGF18-R4-ChIP-F | TCCTTTGGGCTGGTTGAGTG |
| FGF18-R4-ChIP-R | CGTGTGGAAAATGGACCGTG |
| FGF18-R5-ChIP-F | AACCCACAGGTGCTTGTCTC |
| FGF18-R5-ChIP-R | TGGTTTCCTCATTCAAGTCCTCC |
| RT-qPCR primers | |
| GAPDH-F | GCACCGTCAAGGCTGAGAAC |
| GAPDH-R | GGATCTCGCTCCTGGAAGATG |
| FGF18-RT-F | GACGATGTGAGCCGTAAGCA |
| FGF18-RT-R | GAGCTGGGCATACTTGTCCC |
| HBB-RT-F | GAGAACTTCAGGCTCCTGGG |
| HBB-RT-R | GCGAGCTTAGTGATACTTGTGG |
| HBZ-RT-F | CCGGTCAACTTCAAGCTCCT |
| HBZ-RT-R | CTCAGCGGTACTTCTCGGTC |
| HBE1-RT-F | TCACTAGCAAGCTCTCAGGC |
| HBE1-RT-R | AAACAACGAGGAGTCTGCCC |
| SH3PXD2B-RT-F | TGCAGCCTGAAGAAGAGGAG |
| SH3PXD2B-RT-R | CTGGATGACCTCCACCACAG |
| CDKN1A-RT-F | GGCAGACCAGCATGACAGATTT |
| CDKN1A-RT-R | GCTTCCTGTGGGCGGATTA |
| CTCF-RT-F | AATTGAACCTGAGCCAGAGCC |
| CTCF-RT-R | CTGAATGATAGCTGTTGGCTGG |
| SMC1A-RT-F | ACCCAATGGCTCTGGTAAGTC |
| SMC1A-RT-R | AGGTCCCGCAGGGTCTTTA |
| RAD21-RT-F | CATGGTCTTCAGCGTGCTCT |
| RAD21-RT-R | AACTTTGCGGCAGCTTGTTT |
| FXBW11-RT-F | CCAGTGTCAGATGTCTCCAGATAA |
| FXBW11-RT-R | AGTTTCCTTCTGATGGCCTCTT |
| STK10-RT-F | GCAAGGTTTACAAGGCCAAGAA |
| STK10-RT-R | GCTCCTCCTCACTCTTGGTT |

| | |
|---|---|
| NPM1-RT-F | GTTCTCTGGAGCAGCGTTCT |
| NPM1-RT-R | GGCCTTTAGTTCACAACCGAAA |
| UBTD2-RT-F | CACCGGAGTTGCTCTAGGTC |
| UBTD2-RT-R | TCCCTCTTGCTGCGTAGTTG |
| 4C-seq primers | |
| FGF18-Outer-F | TCAGGCCTCCTTGCAAGCTAT |
| FGF18-Outer-R | GAACGAAGCTGCCTAGTATGC |
| FGF18-Nest-F | ACATCGGACACAAGTGCAGAA |
| FGF18-Nest-R | TCCAAAGCTGGCTCGCCTA |
| 3C-PCR primers | |
| FGF18-promoter-3C-F | CAGCCGCAACAATATACCCAGCTCTAAG |
| FGF18-enhancer-3C-R | GCCAGAGAAAAGGTTGGAAGTCATATTCA |
| CRISPR genotyping primers and guide RNAs | |
| CRISPR-step2-F | GCCTTTTGCTGGCCTTTTGCTC |
| CRISPR-step2-R | CGGGCCATTTACCGTAAGTTATGTAACG |
| FGF18-S1-flanking-F | TGGGCTTTTTCTTAAGCTGCC |
| FGF18-S1-flanking-R | TGGCCAAGTTGATTGTGTTAGT |
| FGF18-S1-internal-F | CACCTCAACAAACCGATCACC |
| FGF18-S1-internal-R | GAATCTAATAGCTGAGGACGAGC |
| FGF18-S2-flanking-F | AATCCACTTGAGTTTTTGACACTT |
| FGF18-S2-flanking-R | TCACCACTCAGACACATCAACT |
| FGF18-S2-internal-F | ACTCTACCTGCGACTGTCTTCA |
| FGF18-S2-internal-R | ATCACCACTCAGACACATCAACT |
| FGF18-S1-gRNA-F-F | CACCGTTTTTCCTGTCACGCTGCTC |
| FGF18-S1-gRNA-F-R | AAACGAGCAGCGTGACAGGAAAAAC |
| FGF18-S1-gRNA-S-F | CACCGCTCTGCTTTAAGAGCATCAC |
| FGF18-S1-gRNA-S-R | AAACGTGATGCTCTTAAAGCAGAGC |
| FGF18-S2-gRNA-F-F | CACCGTTGTTTTACGGGCCAGCATA |
| FGF18-S2-gRNA-F-R | AAACTATGCTGGCCCGTAAAACAAC |
| FGF18-S2-gRNA-S-F | CACCGTCTATCTATCTTGTAGAGTC |
| FGF18-S2-gRNA-S-R | AAACGACTCTACAAGATAGATAGAC |

**Supplementary Data**

**Supplementary Data 1**. Hi-C TADs in K562 cells after drug treatment or CRISPR KO (Excel Spreadsheet).

**Supplementary Data 2.** Hi-C loops under different categories in K562 cells after drug treatment or CRISPR KO (Excel Spreadsheet).

**Supplementary Data 3.** Hi-C TADs under different categories in K562 cells after drug treatment (Excel Spreadsheet).

**Supplementary Data 4.** Hi-C loops under different categories in K562 cells after drug treatment (Excel Spreadsheet).

**Supplementary Data 5.** Differentially expressed genes in RNA-seq of K562 cells after drug treatment or CRISPR KO (Excel Spreadsheet). This table listed all the differentially expressed genes from the RNA-seq that we performed for this paper. DESeq2 package was used to analysis differentially expressed genes.

**Supplementary Data 6.** Libraries Used (Excel Spreadsheet). This was a list of all the libraries used in this manuscript.

### Supplementary Figures and Legends

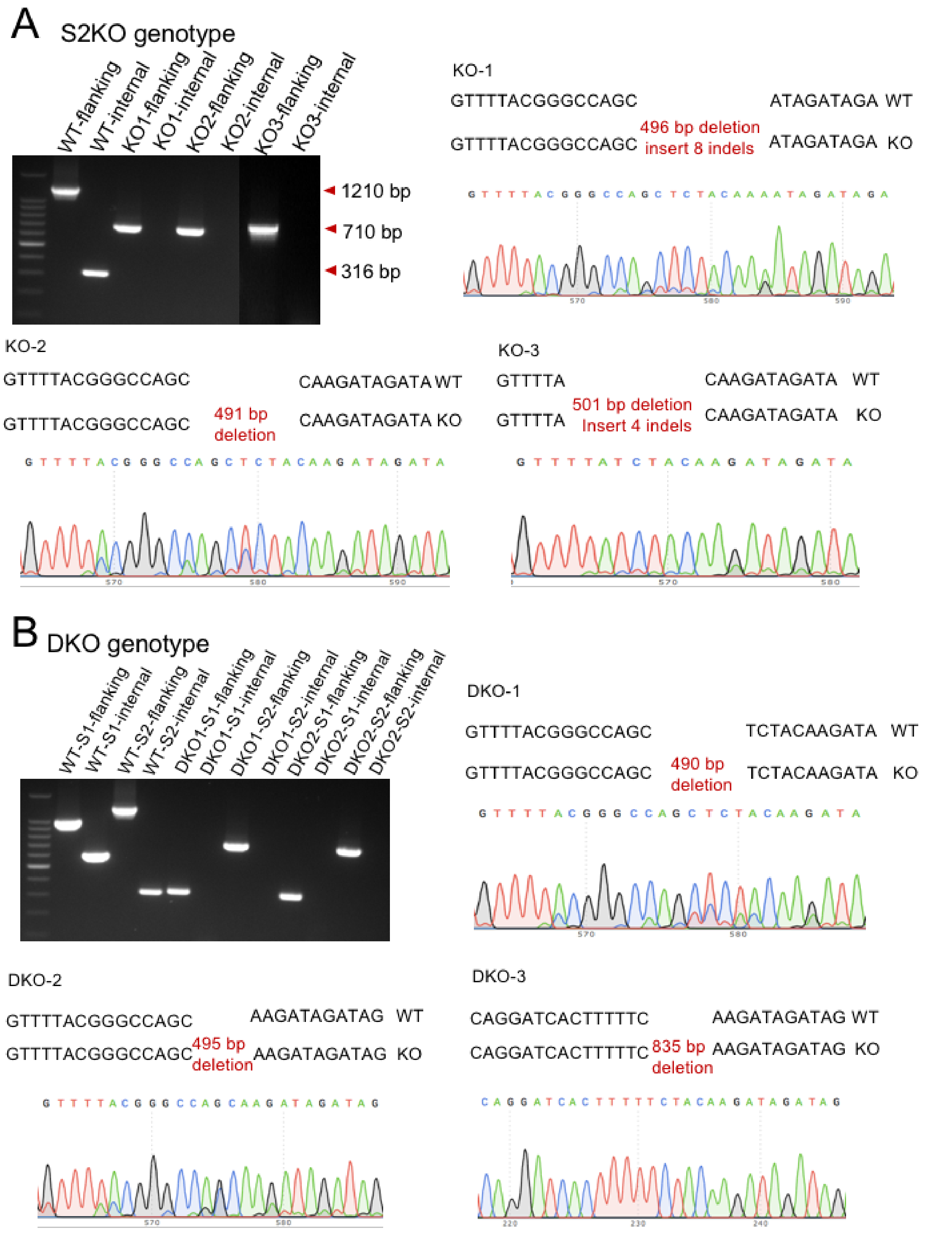

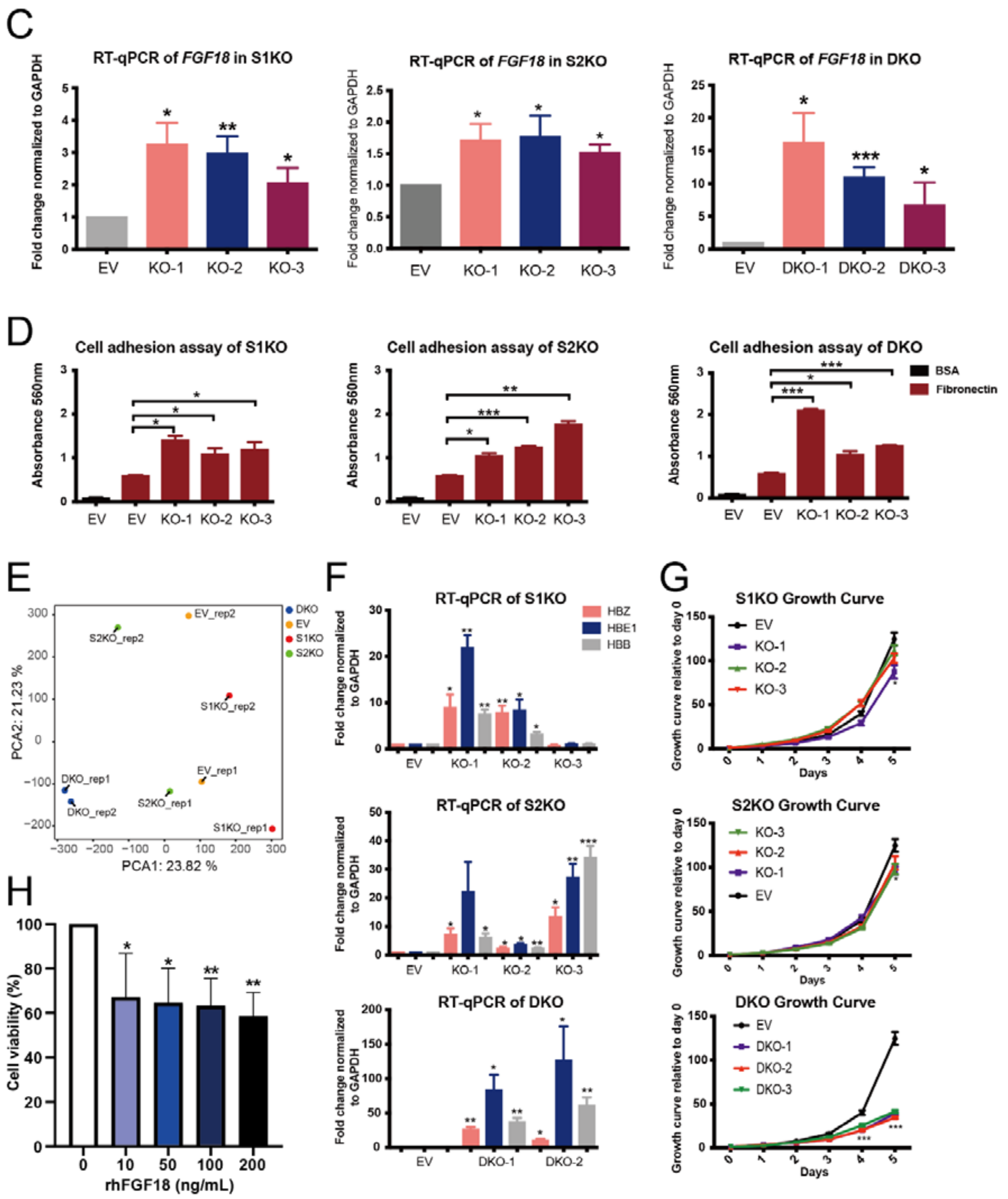

C
RT-qPCR of FGF18 in S1KO
RT-qPCR of FGF18 in S2KO
RT-qPCR of FGF18 in DKO
Fold change normalized to GAPDH
EV
KO-1
KO-2
KO-3
DKO-1
DKO-2
DKO-3
D
Cell adhesion assay of S1KO
Cell adhesion assay of S2KO
Cell adhesion assay of DKO
Absorbance 560nm
BSA
Fibronectin
E
PCA1: 23.82 %
PCA2: 21.23 %
DKO
EV
S1KO
S2KO
EV_rep2
S2KO_rep2
S1KO_rep2
DKO_rep1
EV_rep1
DKO_rep2
S2KO_rep1
S1KO_rep1
F
RT-qPCR of S1KO
RT-qPCR of S2KO
RT-qPCR of DKO
HBZ
HBE1
HBB
Fold change normalized to GAPDH
G
S1KO Growth Curve
S2KO Growth Curve
DKO Growth Curve
Growth curve relative to day 0
Days
H
Cell viability (%)
rhFGF18 (ng/mL)

529

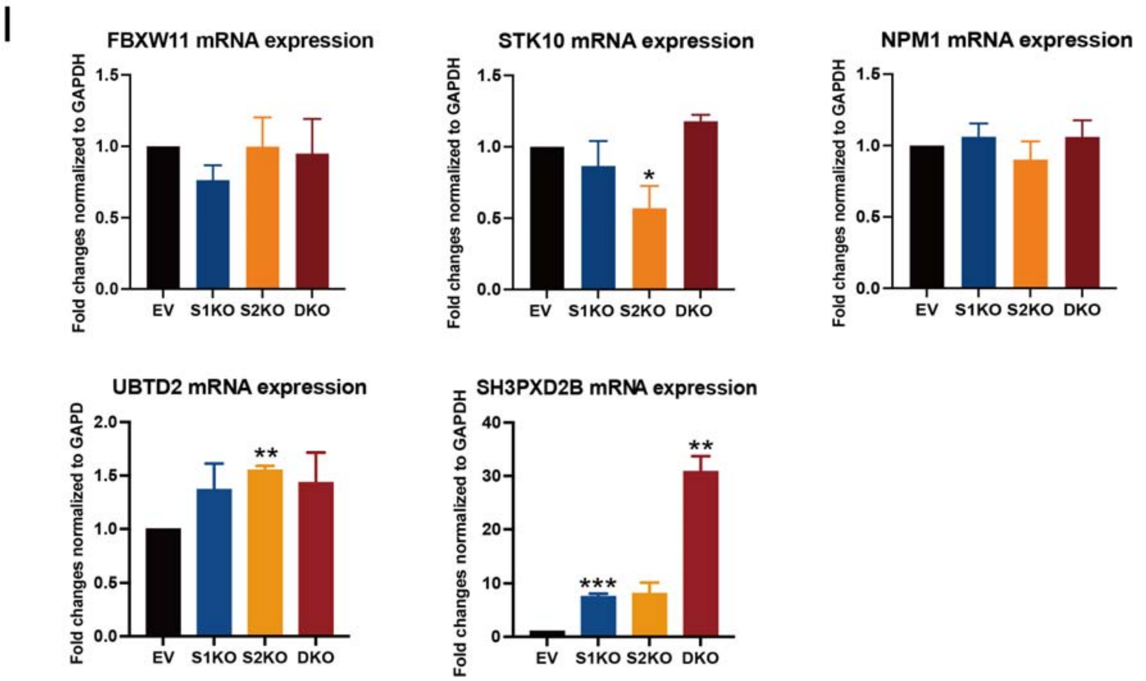

**Figure S1. Two silencers function synergistically to repress *FGF18* expression and control cell identity**. **A & B**. Genotyping of three positive clones of S2KO **(A)** and DKO **(B)** following CRISPR-Cas9 transfection and single cell plate out. Genotyping results of three different clones (KO-1, KO-2, and KO-3) are shown by gel electrophoresis through a pair of flanking and internal primers (top left). Sanger sequencing results of the flanking PCR products for all three knockout clones in both S2KO and DKO conditions. **C**. RT-qPCR analysis of expression of *FGF18* in three CRISPR knockout clones (KO-1, KO-2, and KO-3) and vector control clone ("Empty Vector"; "EV") for S1KO (left), S2KO (middle) and DKO cells (right). Fold change normalized to GAPDH plotted. **D**. Bar graphs showing absorbance values at 560 nm in three CRISPR knockout clones (KO-1, KO-2, and KO-3) and EV for S1KO (left), S2KO (middle) and DKO (right) cells. Fibronectin adhesion assay is performed in S1KO, S2KO and DKO cells by comparing to EV. Bovine Serum Albumin (BSA) used as a negative control. **E**. Principal Components Analysis (PCA) plot of gene expression in EV, S1KO, S2KO and DKO conditions. PCA depicting the variance of the two biological replicates for each condition, as well as the samples from different conditions. Percentage labeled on each axis denotes the extent to which the principal components explained the variation. **F**. RT-qPCR analysis of expression of haemoglobin genes (*HBZ*, *HBE1* and *HBB*) in EV cells, S1KO, S2KO and DKO cells. Fold change normalized to GAPDH plotted. **G**. Growth curves of EV, S1KO, S2KO, and DKO cells. Data calculated as fold change against day 0. **H**. Cell viability of K562 cells either treated with sterile water (control) or recombinant human FGF18 (rhFGF18) at the indicated concentrations for 72 h. Cells were serum starved for 24 h before rhFGF18 treatment. Results are expressed as percentage of cell viability relative to the control. **I**. RT-qPCR analysis of expression of *FGF18* nearby genes (FBXW11, STK10, NPM1, UBTD2 and SH3PXD2B) in EV, S1KO, S2KO and DKO cells. Data shown as average + standard error of mean (SEM). P value calculated by two-tailed student's t-test. P values less than 0.05, 0.01 or 0.001 are denoted as *, ** or ***, respectively.

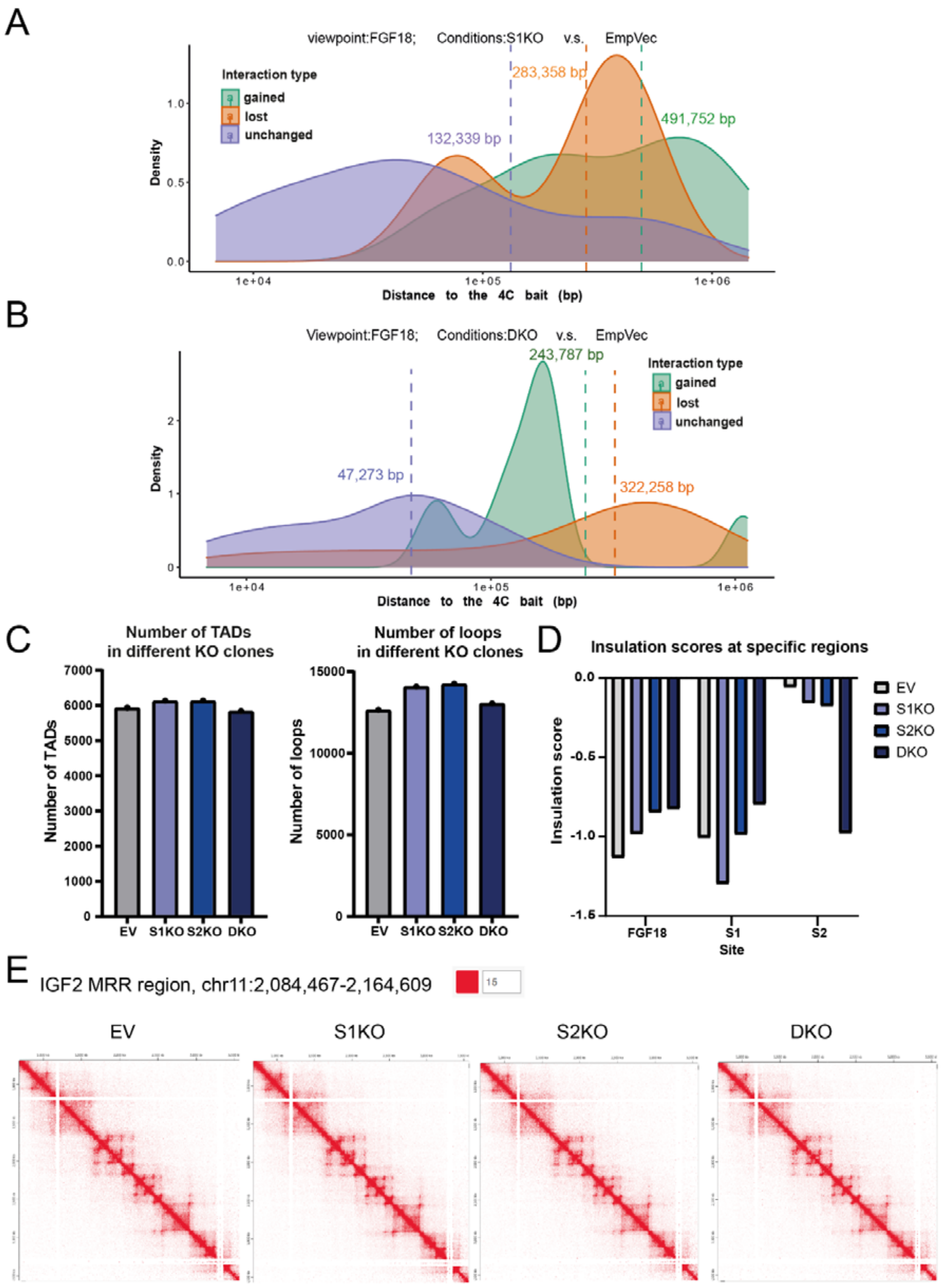
A
viewpoint:FGF18; Conditions:S1KO v.s. EmpVec
Interaction type
gained
lost
unchanged
283,358 bp
491,752 bp
132,339 bp
Density
Distance to the 4C bait (bp)
B
Viewpoint:FGF18; Conditions:DKO v.s. EmpVec
243,787 bp
47,273 bp
322,258 bp
Interaction type
gained
lost
unchanged
Density
Distance to the 4C bait (bp)
C
Number of TADs in different KO clones
Number of TADs
EV S1KO S2KO DKO
Number of loops in different KO clones
Number of loops
EV S1KO S2KO DKO
D
Insulation scores at specific regions
Insulation score
EV
S1KO
S2KO
DKO
FGF18
S1
S2
Site
E
IGF2 MRR region, chr11:2,084,467-2,164,609
15
EV
S1KO
S2KO
DKO

560

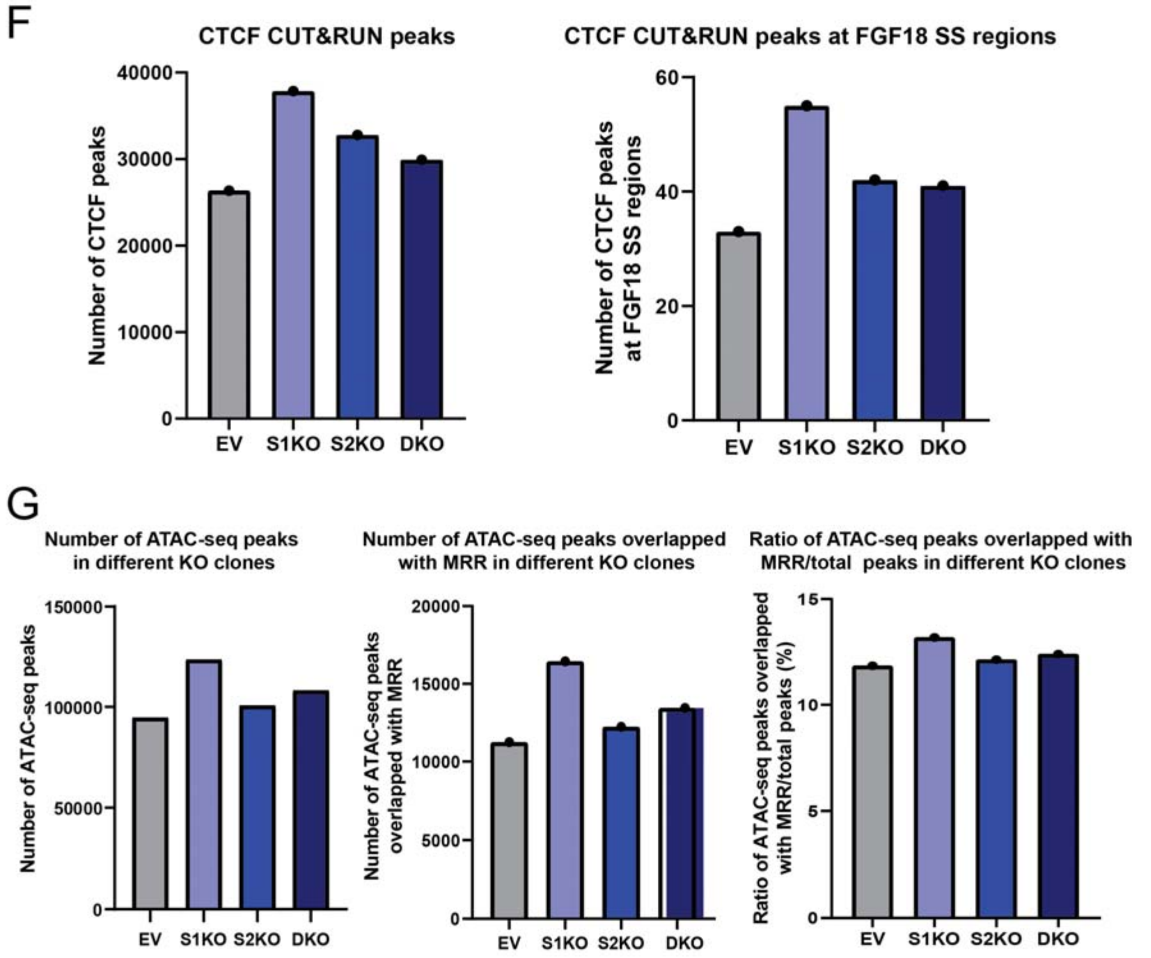

**Figure S2. The changes of TADs and chromatin interactions in different knockout cells**. **A & B**. Density plots of 4C-seq in S1KO vs EV (**A**) and DKO vs EV (**B**) conditions. 4C-seq loops are classified into three different categories (gained, lost and unchanged). Each category is plotted using the distance to the 4C viewpoint. The average distance is shown as a dotted line. **C**. Bar graphs showing number of TADs and loops called from the Hi-C data in EV, S1KO, S2KO and DKO cells. **D**. Bar graph showing detailed insulation scores at the specific regions (*FGF18* gene, S1 and S2) in EV, S1KO, S2KO and DKO cells. **E**. Hi-C matrices at the *IGF2* MRR region in EV, S1KO, S2KO and DKO cells. **F.** Bar graphs showing number of total CTCF Cut & Run peaks and number of CTCF Cut & Run peaks found in FGF18 SS regions in EV, S1KO, S2KO and DKO cells. **G.** Bar graphs showing number of total ATAC-seq peaks, ATAC-seq peaks overlapped with MRR, and ratio of overlapped MRR peaks and total ATAC-seq peaks in EV, S1KO, S2KO and DKO cells.

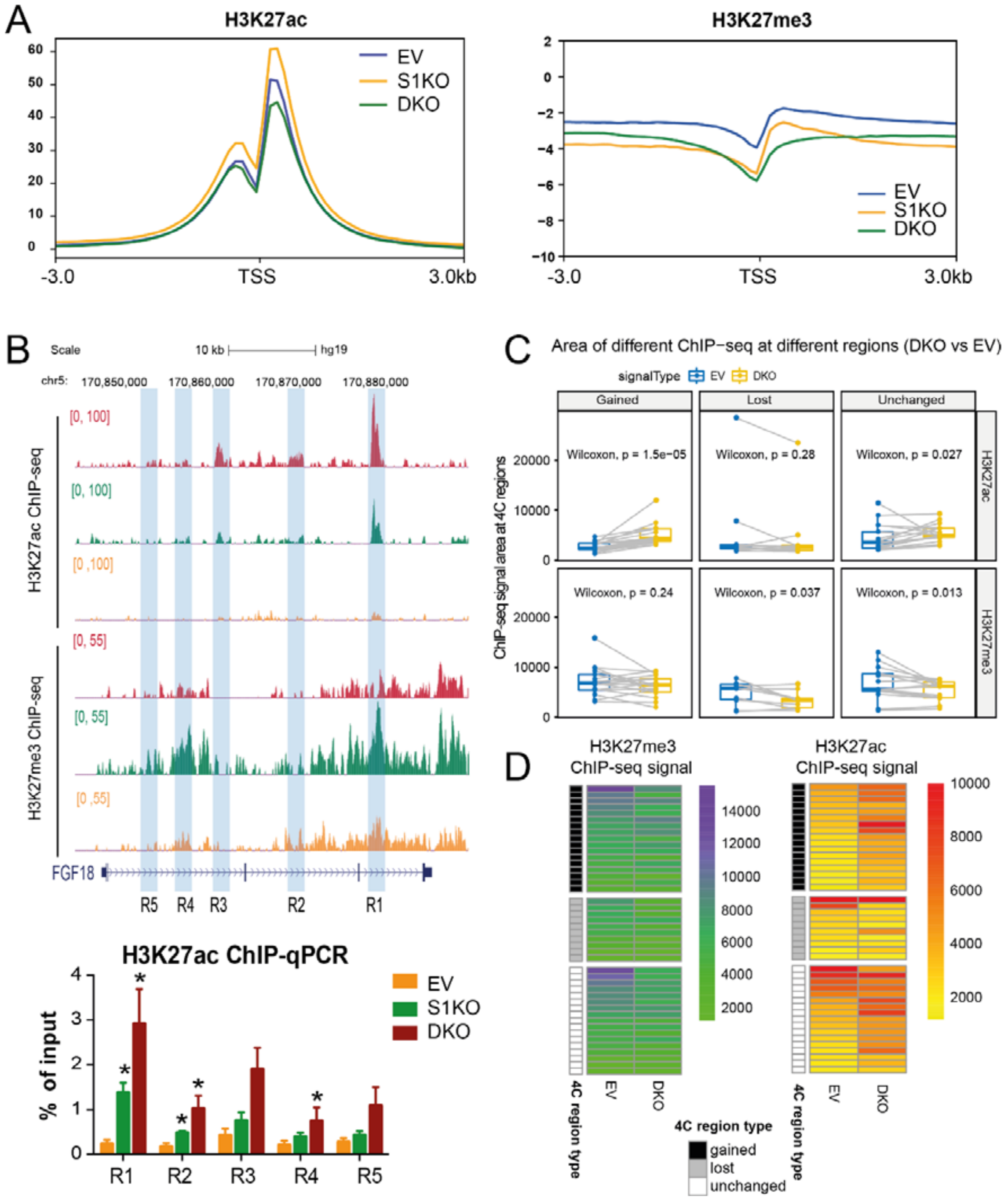
A
H3K27ac
EV
S1KO
DKO
-3.0
TSS
3.0kb
H3K27me3
B
Scale
10 kb
hg19
chr5: 170,850,000 170,860,000 170,870,000 170,880,000
H3K27ac ChIP-seq
H3K27me3 ChIP-seq
FGF18
R5 R4 R3 R2 R1
H3K27ac ChIP-qPCR
% of input
C
Area of different ChIP−seq at different regions (DKO vs EV)
signalType EV DKO
Gained
Lost
Unchanged
ChIP−seq signal area at 4C regions
Wilcoxon, p = 1.5e−05
Wilcoxon, p = 0.28
Wilcoxon, p = 0.027
Wilcoxon, p = 0.24
Wilcoxon, p = 0.037
Wilcoxon, p = 0.013
H3K27ac
H3K27me3
D
H3K27me3 ChIP-seq signal
H3K27ac ChIP-seq signal
4C region type
4C region type
gained
lost
unchanged

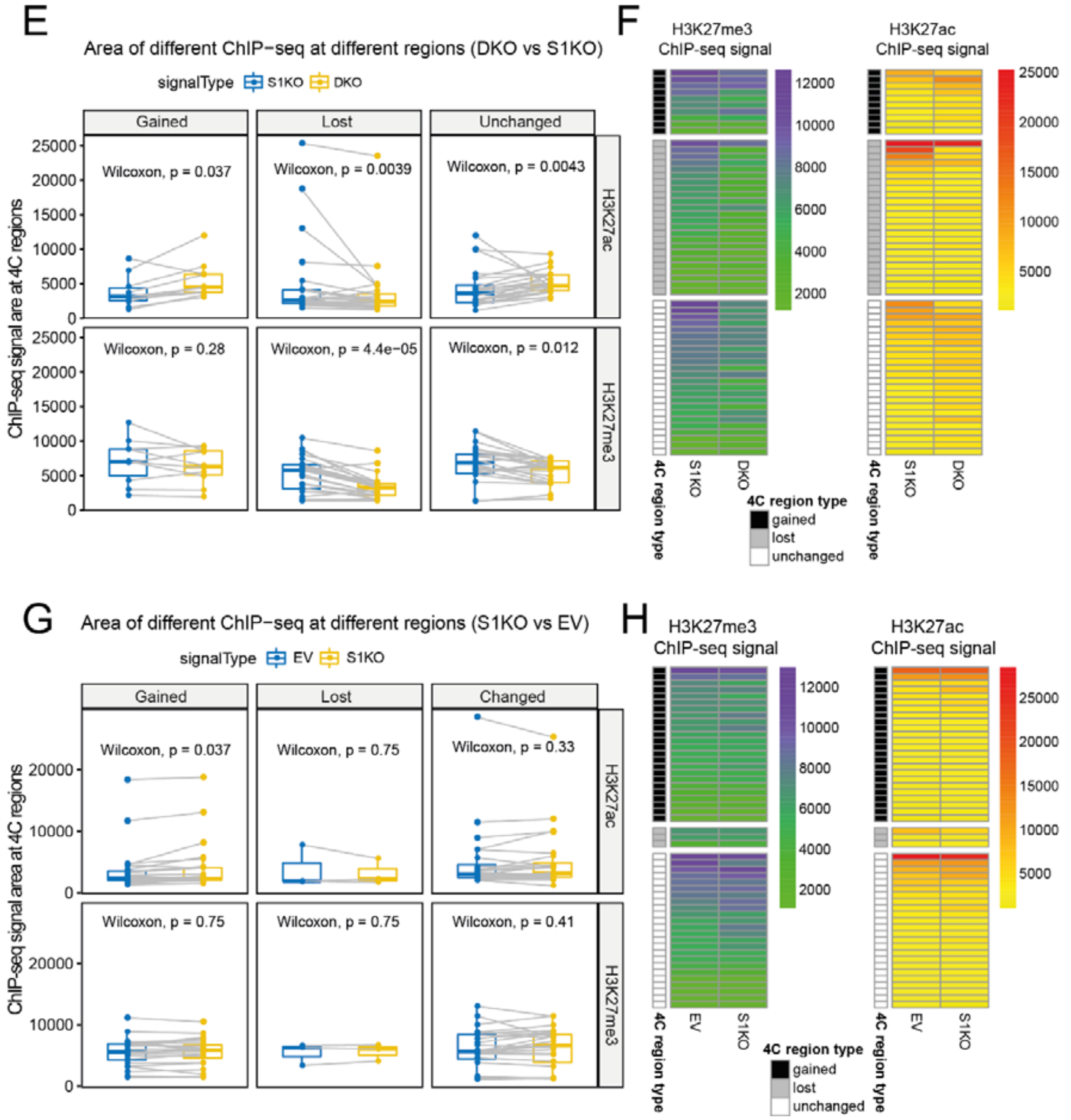
E
Area of different ChIP−seq at different regions (DKO vs S1KO)
signalType S1KO DKO
Gained
Lost
Unchanged
H3K27ac
H3K27me3
ChIP-seq signal are at 4C regions
Wilcoxon, p = 0.037
Wilcoxon, p = 0.0039
Wilcoxon, p = 0.0043
Wilcoxon, p = 0.28
Wilcoxon, p = 4.4e−05
Wilcoxon, p = 0.012
F
H3K27me3 ChIP-seq signal
H3K27ac ChIP-seq signal
4C region type
S1KO
DKO
4C region type
gained
lost
unchanged
G
Area of different ChIP−seq at different regions (S1KO vs EV)
signalType EV S1KO
Gained
Lost
Changed
H3K27ac
H3K27me3
ChIP-seq signal area at 4C regions
Wilcoxon, p = 0.037
Wilcoxon, p = 0.75
Wilcoxon, p = 0.33
Wilcoxon, p = 0.75
Wilcoxon, p = 0.75
Wilcoxon, p = 0.41
H
H3K27me3 ChIP-seq signal
H3K27ac ChIP-seq signal
4C region type
EV
S1KO
4C region type
gained
lost
unchanged

576

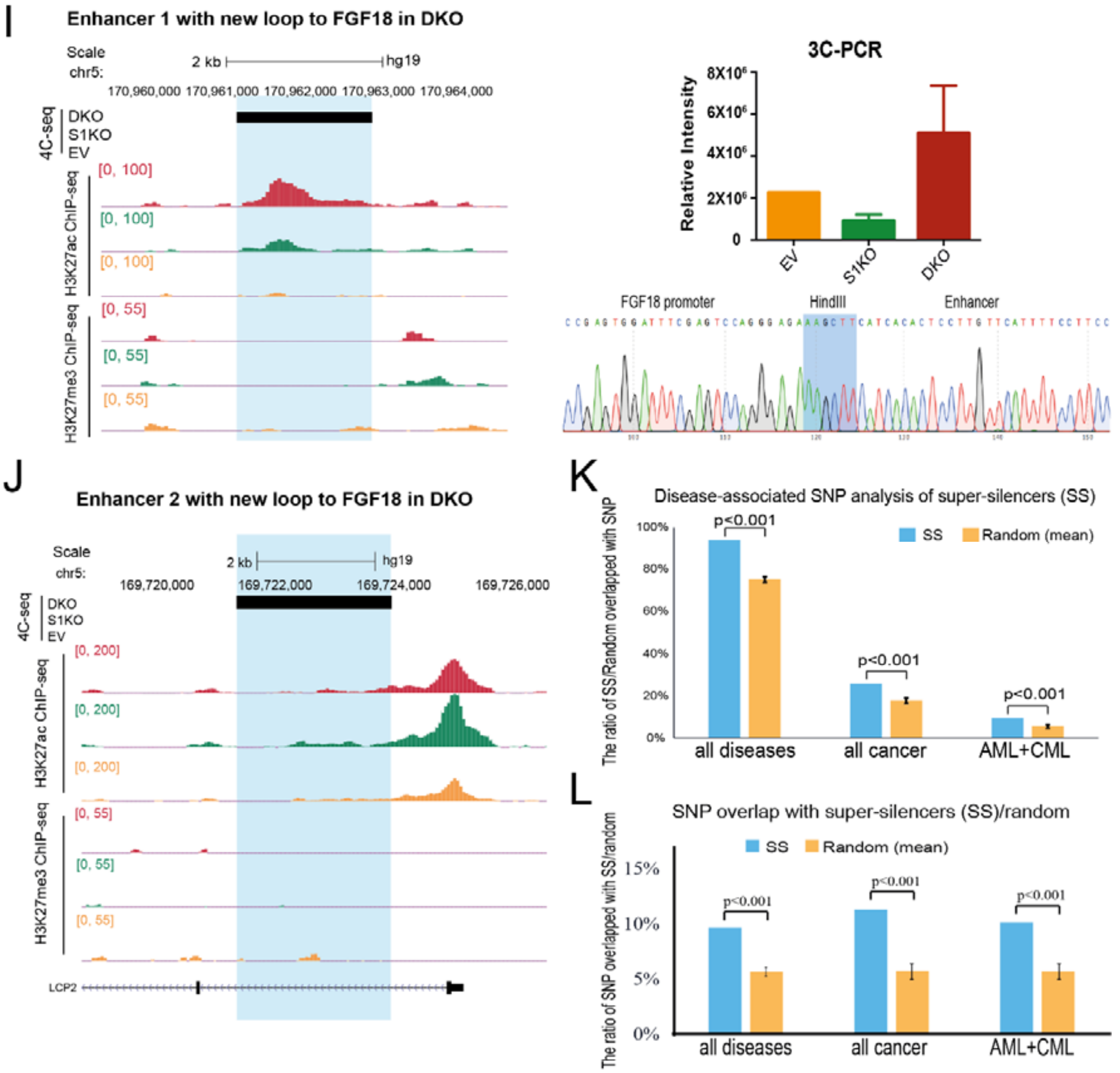

**Figure S3. The unchanged loops and gained loops at *FGF18* locus in DKO cells demonstrate increased H3K27ac signals and decreased H3K27me3 signals**. **A**. Mean plots showing average distribution of H3K27ac ChIP-seq signals (left) and H3K27me3 ChIP-seq signals (right) around the transcription start site (TSS) of known genes in EV, S1KO, and DKO conditions. X-axis represents genome distance to TSS of genes. **B**. Screenshot showing H3K27ac ChIP-seq and H3K27me3 ChIP-seq signal tracks at *FGF18* gene region in EV, S1KO and DKO cells. Bar graph showing ChIP-qPCR against H3K27ac marks is performed at five regions along *FGF18* gene body (R1-R5). Data shown as % of input. Three different replicates are performed for ChIP-seq and ChIP-qPCR. **C and D**. Boxplots (**C**) and heatmaps (**D**) showing ChIP-seq signal changes of H3K27me3 and H3K27ac at different types of 4C regions (gained, lost and unchanged 4C loops) in EV and DKO cells. The same 4C regions are connected by gray lines in the boxplot. P values indicated in each boxplot. **E and F**. Boxplots (**E**) and heatmaps (**F**) of ChIP-seq signal changes of H3K27me3 and H3K27ac at different types of 4C regions (gained, lost and unchanged 4C loops) in S1KO and DKO cells. The same 4C regions are connected by gray lines in the boxplot. P values indicated in each boxplot. **G and H**. Boxplots (**G**) and heatmaps (**H**) showing ChIP-seq signal changes of H3K27me3 and H3K27ac at different types of 4C regions (gained, lost and unchanged 4C loops) in EV and S1KO cells. The same 4C regions are connected by gray lines in the boxplot. P values indicated in each boxplot. **I**. Screenshot depicting aligned H3K27ac ChIP-seq, H3K27me3 ChIP-seq, and 4C-seq signal tracks of enhancer 1 (one of the new enhancers found in DKO cells) in EV, S1KO and DKO cells. Enhancer 1 is highlighted in blue. 3C-PCR of enhancer 1 in EV, S1KO and DKO cells by two independent 3C libraries. Data are relative intensity measured with ImageJ. The panel below are the sanger sequencing results for the 3C ligated fragment. *FGF18* promoter sequence, HindIII cut site and enhancer 1 sequence are indicated. **J**. Screenshot depicting aligned H3K27ac ChIP-seq, H3K27me3 ChIP-seq and 4C-seq signal tracks of enhancer 2 (another new enhancer found in DKO cells) in EV, S1KO and DKO cells. Enhancer 2 is highlighted in blue. **K and L**. Bar graphs showing ratio of SS or random regions overlapped with SNPs (**K**) or ratio of SNPs overlapped with SS or random regions (**L**) in different SNP groups (all diseases, all cancer, and AML + CML). Disease-associated SNPs were downloaded from GWAS [28]. P values comparing super-silencers with random regions for each group were indicated. P value calculated by two-tailed student's t-test. P value less than 0.05 shown as *.

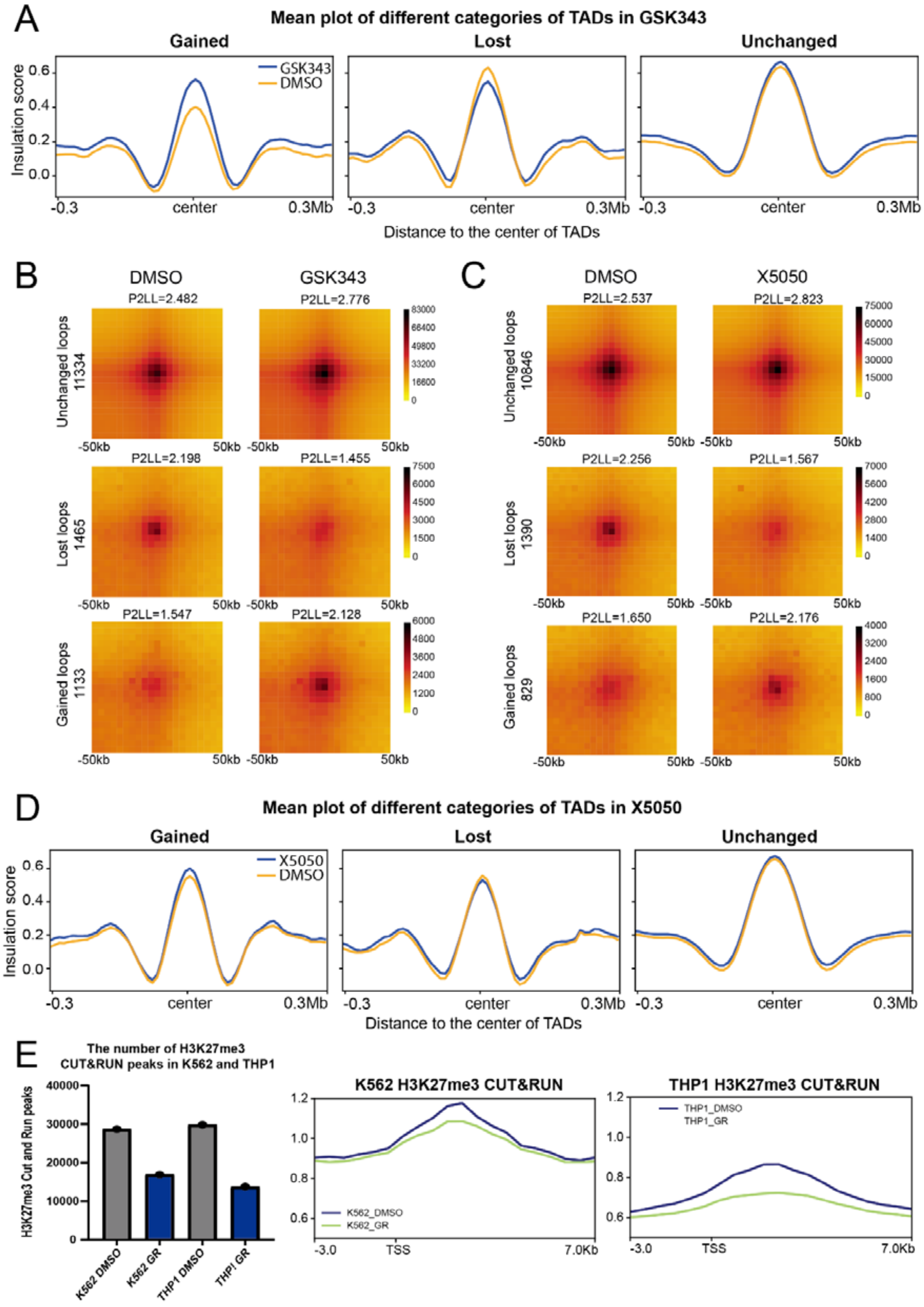
A
Mean plot of different categories of TADs in GSK343
Gained
Lost
Unchanged
GSK343
DMSO
Insulation score
Distance to the center of TADs
B
DMSO
GSK343
P2LL=2.482
P2LL=2.776
Unchanged loops 11334
P2LL=2.198
P2LL=1.455
Lost loops 1465
P2LL=1.547
P2LL=2.128
Gained loops 1133
C
DMSO
X5050
P2LL=2.537
P2LL=2.823
Unchanged loops 10846
P2LL=2.256
P2LL=1.567
Lost loops 1390
P2LL=1.650
P2LL=2.176
Gained loops 829
D
Mean plot of different categories of TADs in X5050
Gained
Lost
Unchanged
X5050
DMSO
Insulation score
Distance to the center of TADs
E
The number of H3K27me3 CUT&RUN peaks in K562 and THP1
H3K27me3 Cut and Run peaks
K562 DMSO
K562 GR
THP1 DMSO
THP1 GR
K562 H3K27me3 CUT&RUN
K562_DMSO
K562_GR
THP1 H3K27me3 CUT&RUN
THP1_DMSO
THP1_GR

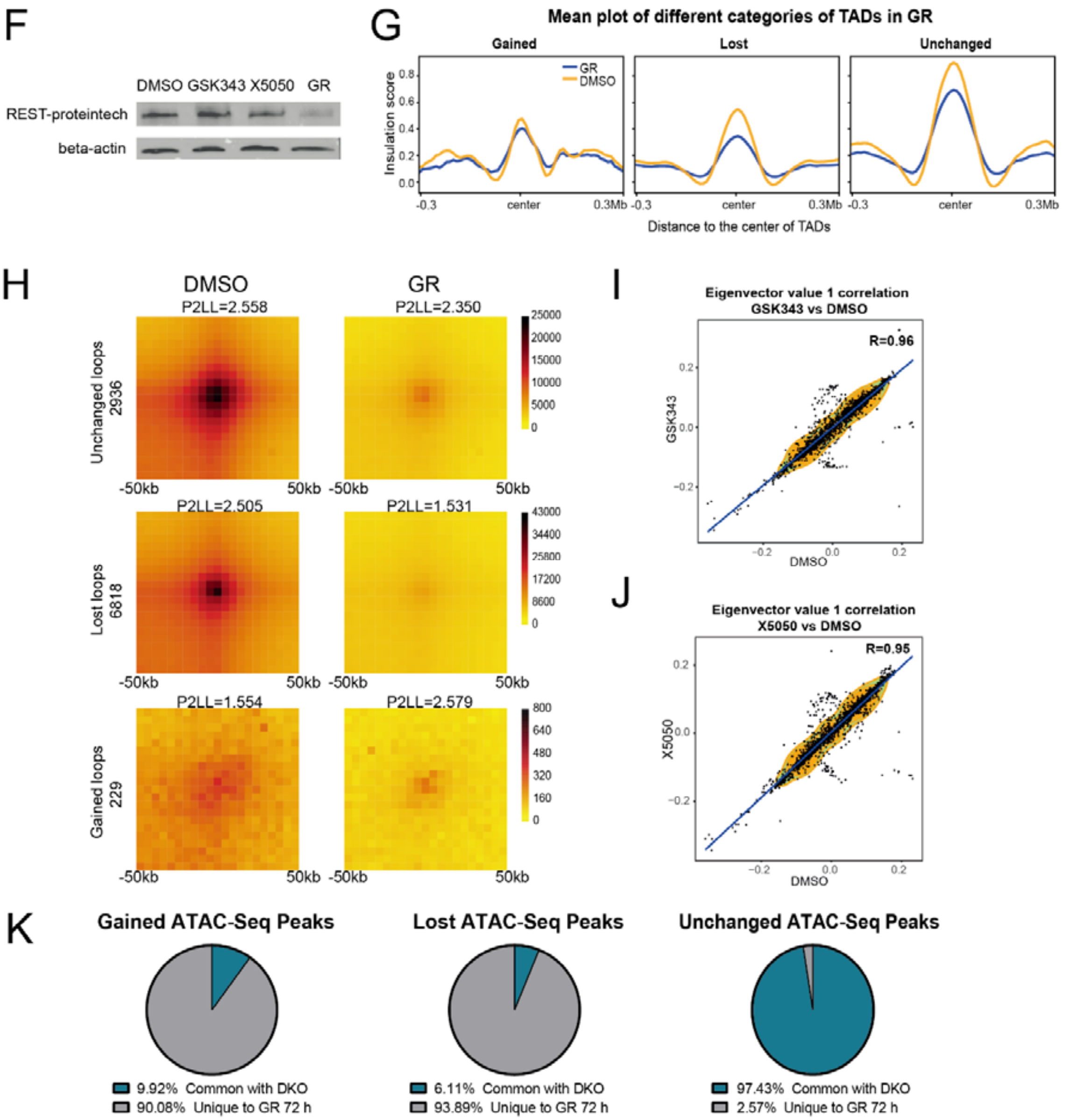

**Figure S4. Combinational treatment of GSK343 and X5050 in K562 cells leads to synergistic losses of TADs and loops. A**. Mean plots describing genome-wide insulation score around TADs in the unchanged, lost, and gained TADs in GSK343 vs DMSO condition. **B and C**. Aggregate peak analysis (APA) for unchanged, lost and gained loops in GSK343 vs DMSO (**B**) and X5050 vs DMSO conditions (**C**). Loops aggregated at the center of a 50kb window at 5kb resolution. The ratios of signal at the peak signal enrichment (P) to the average signal at the lower left corner of the plot (LL) (P2LL) are indicated to show the normalized intensity of all loops. **D**. Mean plots describing genome-wide insulation score around TADs in the unchanged, lost and gained TADs in X5050 vs DMSO condition. **E**. Bar graph showing number of total H3K27me3 Cut & Run peaks in K562 and THP1 cells treated with DMSO or GR for 24 h (left). Mean plots showing global H3K27me3 levels around TSS in K562 cells (middle) and THP1 cells (right) treated with DMSO or GR for 24 h. **F**. Western blot showing REST protein levels in K562 cells following DMSO, GSK343, X5050 or GR treatments for 72 h. **G**. Mean plots depicting genome wide insulation score around TADs in the unchanged, lost and gained TADs in GR vs DMSO condition. **H**. APA for the unchanged, lost and gained loops in GR vs DMSO condition. Loops aggregated at the center of a 50kb window at 5kb resolution. The ratios of signal at the peak signal enrichment (P) to the average signal at the lower left corner of the plot (LL) (P2LL) are indicated to show the normalized intensity of all loops. **I and J**. Density plots depicting global correlation between the eigenvector value in GSK343 vs DMSO (**I**) and X5050 vs DMSO conditions (**J**) at 1Mb resolution. X-axis represents eigenvector value in the DMSO condition, while Y-axis represents eigenvector value in GSK343 or X5050 treatment conditions at the same locus. **K**. Pie charts depicting distribution of gained, unchanged and lost peaks that are common and unique in GR-treated cells (72 h) as compared to DKO cells.

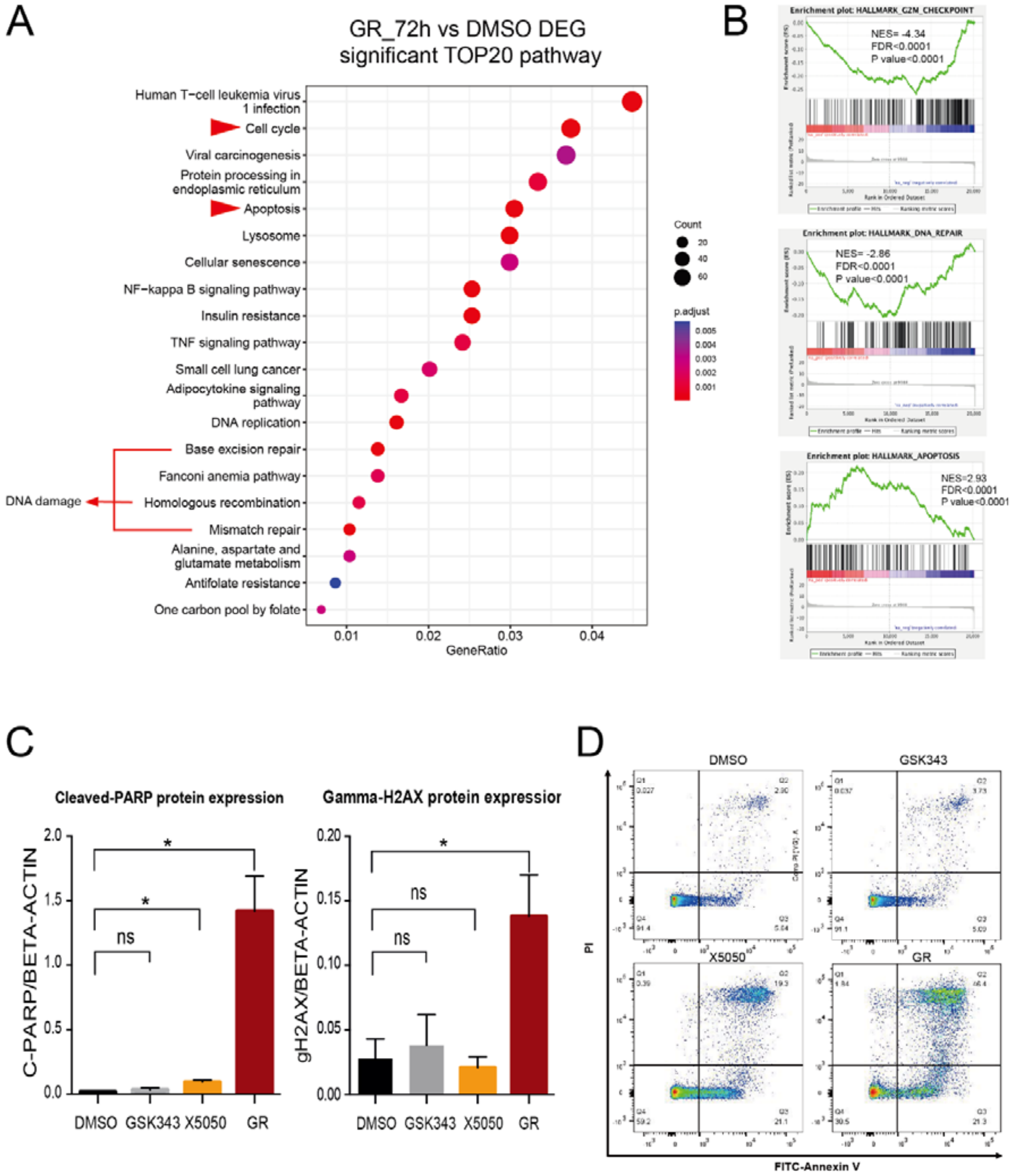

**Figure S5. GR treatment leads to apoptosis and cell cycle arrest through upregulation of super-silencer controlled genes.** Graph showing top 20 significant KEGG pathways based on Differentially Expressed Genes (DEGs) in GR at 72 h versus DMSO RNA-seq. X-axis shows percentage of DEGs in each pathway, while Y-axis shows names of enriched KEGG pathways. Pathways are ranked in order of significance, with the most significant pathway at the top. **B**. Enrichment plots showing gene sets related to G2/M checkpoint, DNA repair and apoptosis in GR vs DMSO. Normalized enrichment score (NES), false recovery rates (FDR) and p value are indicated for each gene set. **C**. Bar graphs showing ratios of cleaved-PARP (left) and gamma-H2AX (right) protein levels normalized to beta-actin protein levels in K562 cells at 72 h following DMSO, GSK343, X5050 or GR treatments. **D**. Scatter plots of Annexin V (AV) and propidium iodide (PI) staining for K562 cells treated with DMSO, GSK343,

X5050 or GR for 72 h. Results shown are representative of three independent experiments. Quadrant 1, necrotic cells AV-/PI+; Quadrant 2, late apoptotic cells AV+/PI+; Quadrant 3, early apoptotic cells AV+/PI-; Quadrant 4, living cells AV-/PI-. Quantitative analysis of the percentage of total apoptotic cells (early apoptotic cells and late apoptotic cells) by flow cytometry are shown. Data shown as average + SEM. P value calculated by two-tailed student's t-test. P value less than 0.05, 0.01, 0.001 or 0.0001 is shown as *, **, *** or ****, respectively.

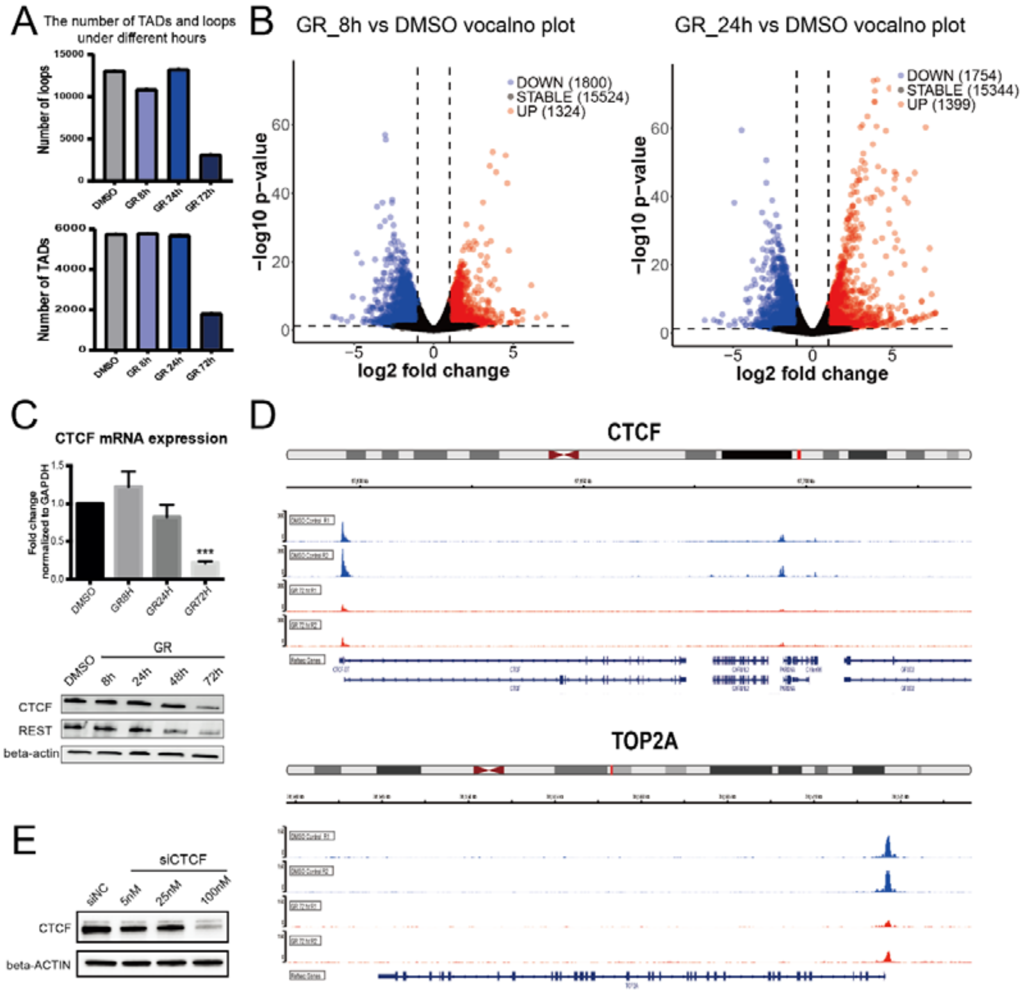

**Figure S6. Gradual depletion of CTCF and TOP2A explains the kinetics and amplitude of the loss of TADs and loops caused by GR treatment. A**. Bar graphs showing number of TADs and loops called from the Hi-C data in DMSO, GR_8 h, GR_24 h and GR_72h conditions. **B**. Volcano plots depicting DEGs in K562 cells following GR treatment for 8 h or 24 h. The number of downregulated, stable and upregulated genes is indicated. **C**. RT-qPCR analysis of expression of *CTCF* (top) and protein levels of CTCF and REST (bottom) in K562 cells treated with DMSO or GR for the indicated time points. **D**. Screenshots of ATAC-seq signal tracks at the promoters and gene body regions of CTCF (top) and TOP2A (bottom) in K562 cells treated with DMSO or GR for 72 h. **E**. Western blot showing CTCF protein levels in K562 cells transfected with siScramble or siCTCF. Data shown as average + SEM. P value calculated by two-tailed student's t-test. P value less than 0.05, 0.01 or 0.001 shown as *, ** or ***, respectively.

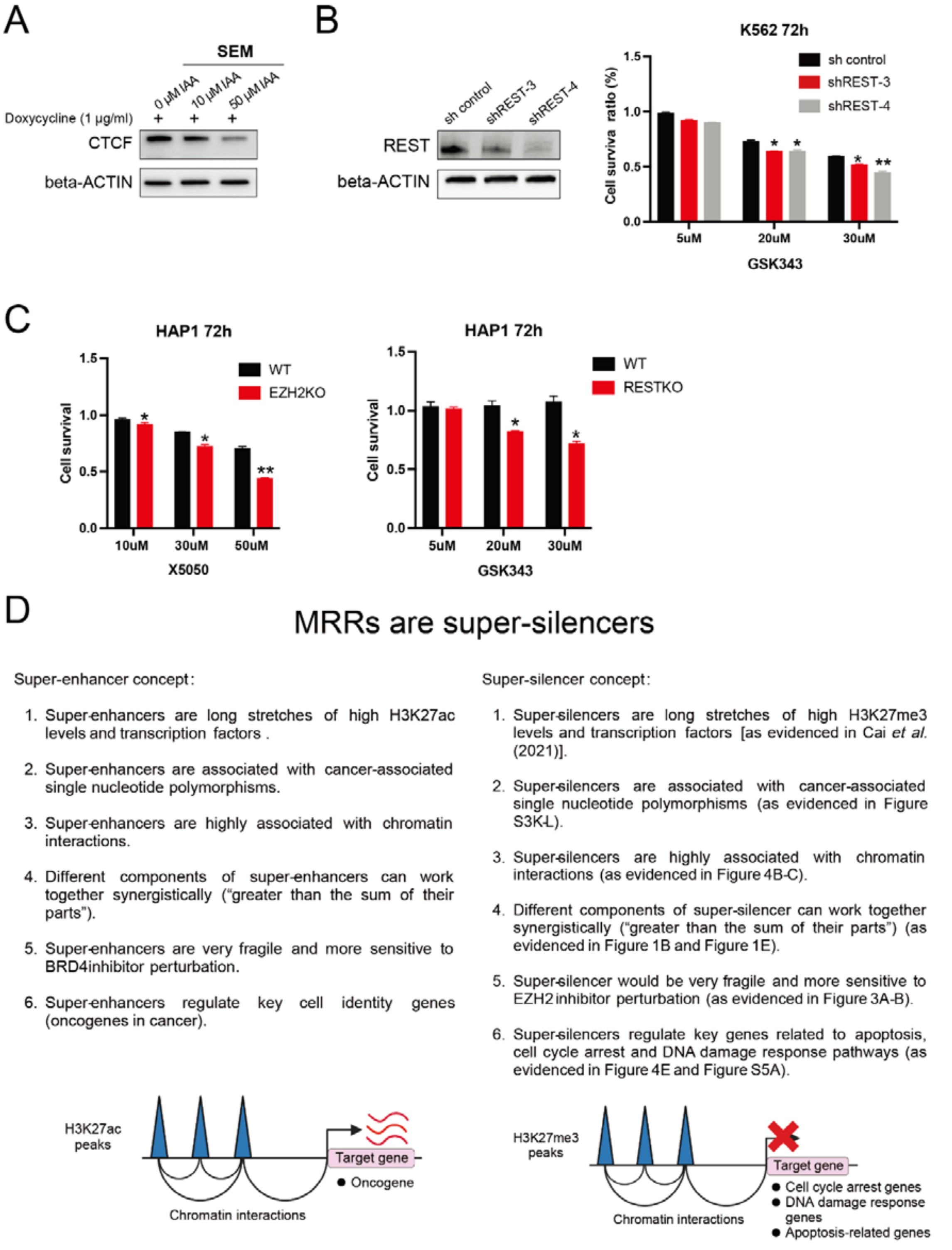

**Figure S7. GR exerts synergistic antitumor effects. A**. Western blot showing CTCF and β-actin protein levels in SEM cells either treated with or without IAA for 48 h. **B**. Western blot showing REST protein levels in two REST knockdown clones (shREST-3 and shREST-4) (left). Bar graph showing percentage of cell viability in REST-depleted K562 cells treated with increasing concentrations of GSK343 (right). **C**. Bar graphs showing percentage of cell viability in EZH2 knockout HAP1 cells (left) and REST knockout HAP1 cells (right) treated with increasing concentrations of X5050 or GSK343, respectively. **D**. Schematics describing concepts of super-enhancer (left) and super-silencer (right). Data shown as average + SEM. P value calculated by two-tailed student's t-test. P value less than 0.05, 0.01 or 0.001 shown as *, ** or ***, respectively.
